# Transposable Element insertions Repurpose Immune Endonucleases as Drivers of Reproductive Isolation in Mice

**DOI:** 10.64898/2026.09.24.753999

**Authors:** Tabitha Rucker, Sandrine Vandormael-Pournin, Joris Mordier, Lolita Lecompte, Aurélie Tessandier, Pablo Navarro, Deborah Bourc’his, Antoine Molaro, Michel Cohen-Tannoudji

**Affiliations:** Institut Pasteur, Université Paris Cité, CNRS UMR3738, Epigenomics, Proliferation, and the Identity of Cells, Department of Developmental and Stem Cell Biology, F-75015 Paris, France; Institut Pasteur, Université Paris Cité, CNRS UMR3738, Early Mammalian Development and Stem Cell Biology, F-75015, Paris, France; Institute of Genetics, Reproduction & Development, CNRS UMR6293, INSERM U.1103, Université Clermont Auvergne, 28 Place Henri Dunant, Clermont-Ferrand, France; INSERM U934, CNRS UMR3215, Institut Curie, PSL Research University, Paris, F-75005 Paris, France

**Author notes:** These authors contributed equally to the work (Tabitha Rucker, Sandrine Vandormael-Pournin).

## Abstract

The emergence of reproductive barriers between closely related populations is a fundamental step in speciation, yet the underlying genetic mechanisms remain poorly understood in mammals. Here we resolve the DDK syndrome, a long-standing model of hybrid incompatibility in mice, by showing that two independent retrotransposon insertions rewire immune *Schlafen* endoribonuclease genes expression, causing hybrid embryo death. A DDK-private MERVL element drives ectopic oocyte expression of *Slfn1*, while an intragenic ETn/MusD insertion in incompatible strains is associated with zygotic *Slfn15* expression. SLFN1-SLFN15 heterodimerization constitutively activates tRNA ribonuclease activity, leading to global translation inhibition, and integrated stress response activation. Abolishing SLFN1 catalytic activity or SLFN15 expression rescues syndromic embryo development. Our findings uncover a previously unrecognized speciation mechanism in which transposable elements repurpose immune effectors as drivers of postzygotic reproductive isolation.

## INTRODUCTION

Understanding the genetic basis of speciation has been a central question in evolutionary biology for decades. In sexually reproducing organisms, current speciation theories predict that in the absence of gene flow, independently evolving populations will eventually accumulate genetic differences that cause inviability or sterility upon crossing^1–3^. Although such reproductive barriers have been documented across the tree of life, their underlying genetic basis has been resolved in only a handful of species, predominantly invertebrates and plants^4,5^. Many incompatibility genes arise from antagonistic co-evolution or genetic conflicts, leading to repeated positive selection of diverging alleles between closely related populations^6–8^. Several conflicts have been proposed to contribute to reproductive isolation, including host/pathogen interactions, the suppression of germline selfish genetic elements (e.g., meiotic drivers and transposable elements), and parental-effect genes influencing zygotic development^3,7,8^. However, the evolutionary and molecular mechanisms underlying reproductive isolation remain largely unknown in mammals, limiting our understanding of how speciation occurs in this lineage.

To date, the only speciation gene identified in mammals is *Prdm9*, which licenses sites of programmed double-strand breaks during meiosis. PRDM9 zinc fingers undergo rapid intraspecies adaptation as a result of inevitable biased gene conversion during break repair and, as such, *Prdm9* was originally identified as the primary locus responsible for hybrid male sterility between *Mus musculus (M. m.) musculus* and *M. m. domesticus* subspecies^9,10^. Since PRDM9 molecular function is conserved beyond rodents, it is thought to drive cryptic hybrid incompatibility across mammals^11,12^.

The other emblematic yet unresolved model of mammalian reproductive isolation is the DDK syndrome^13,14^. This hybrid incompatibility phenotype occurs when females from the DDK Japanese laboratory mouse inbred strain are mated with males from other *M. m. domesticus* laboratory strains such as BALB/c or C57BL/6^15–17^ (referred to as incompatible strains, “Incomp”, Figure 1A). These crosses result in the death of nearly all F1 embryos at the morula-to-blastocyst transition, while reciprocal crosses involving a DDK male are fully fertile (Figure 1A). From a speciation perspective, some wild-derived *M. m. domesticus* inbred strains (e.g. WSB and PERC but not LEWES or PERA), as well as strains from other *M. m.* subspecies (e.g. PWK (*M. m. musculus*), MOLC (*M. m. molosinus*), and CAST (*M. m. castaneus*)) produce viable offspring when crossed with DDK females (Figure 1A, compatible lines, “Comp”)^18^. This pattern suggests that the incompatible paternal allele arose relatively recently during *M. m. domesticus* lineage expansion, less than 0.5 Mya.

**Figure 1:**
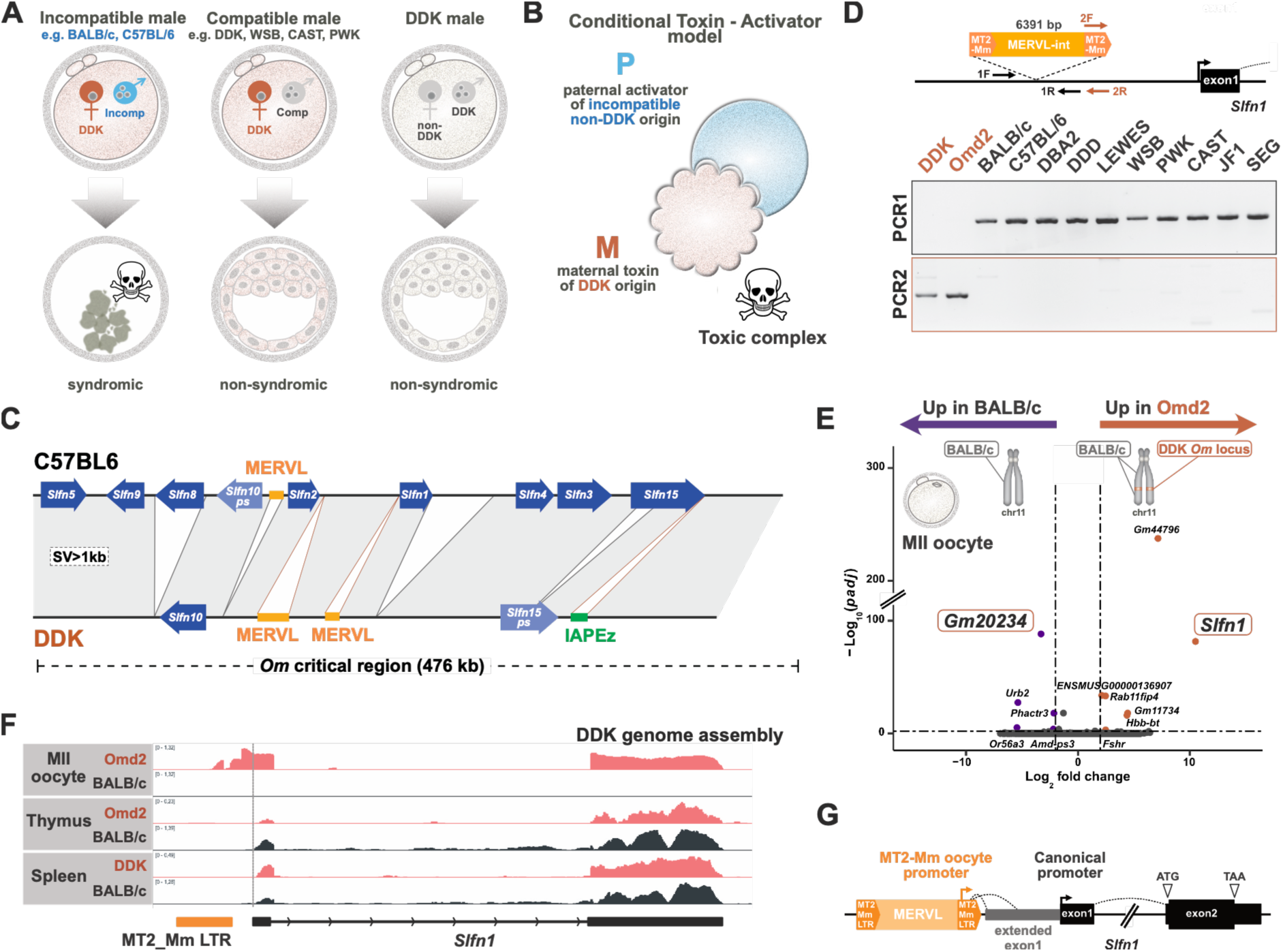
A DDK-private full-length MERVL insertion drives ectopic *Slfn1* expression in oocytes. **A.** The DDK syndrome, an early embryonic lethality occurring at the morula-to-blastocyst transition, is observed exclusively when DDK females are mated with males from incompatible strains. Reciprocal crosses, as well as crosses involving DDK males are fully fertile. **B.** Conditional Toxin-Activator model proposing that a toxic complex can be assembled only in (DDK x incompatible)F1 embryos through the interaction of a maternally deposited DDK-specific factor with an incompatible paternally expressed factor. **C.** Schematic comparison of the *Ovum mutant* critical region between the DDK genome assembly and the mm39 C57BL/6 reference genome (incompatible strain), highlighting structural variants (SVs) larger than 1 kb. No SVs exceeding 1kb were identified in the portion of the *Om* region not depicted in the schematic. **D.** Genomic PCR using primers flanking the MERVL full-length integration site upstream of *Slfn1* (PCR1) or specific to the MERVL insertion (PCR2). **E.** Volcano plot showing differentially expressed genes (FDR<0.01; |log2(FC)| > 2) between BALB/c and Omd2 MII oocytes. Genes located within the *Om* critical region are circled in red. **F.** IGV plots showing *Slfn1* read coverage after alignment to the DDK genome assembly. **G.** Schematic representation of the gene structure of canonical and oocyte-specific *Slfn1* transcript isoforms.

Based on previous genetic, molecular and experimental embryology studies^13,14,17,19,20^, we propose a Conditional Toxin - Activator model whereby a maternally deposited DDK oocyte product compromises development only upon interaction with an incompatible paternal product (Figure 1B). Both the maternal and paternal factors have been mapped to a 476 kb critical interval on the distal region of chromosome 11, termed *Ovum mutant (Om)* ^18,21,22^. Remarkably, 9 of the 12 protein-coding genes within *Om* belong to the *Schlafen* (*Slfn*) family of RNases. Extensive phylogenetic studies in mammals have shown that *Slfn* genes undergo rapid evolutionary diversification, recurrent gene conversion, extensive copy number variation, and strong positive selection affecting their functional domains and their dimerization interfaces^23,24^. Originally identified as regulators of immune cell proliferation and differentiation^25,26^, SLFN proteins have since emerged as key effectors of innate immune and antiviral responses^25–27^. They share a conserved N-terminal “Schlafen core domain” with RNA binding and cleavage activity and, in longer family members, an additional C-terminal helicase-like extension with additional nucleotide binding or regulatory functions^28^. Mechanistically, SLFN proteins primarily function as endoribonucleases that specifically cleave transfer RNAs (tRNAs), leading to codon-specific ribosome stalling, global translational repression, and subsequent activation of the integrated stress response^29–33^. Besides their well-established roles in immunity, no reproductive function has been attributed to *Slfn* genes to date.

## RESULTS

### The DDK strain has a private MERVL full-length integrant upstream of *Slfn1*

To identify genetic variations responsible for the incompatibility phenotype, we generated a full-length, draft *de novo* assembly of the DDK mouse genome (Figure S1A-C). Comparing DDK to the reference C57BL/6 mouse genome identified ∼5.8 million single nucleotide variants (SNVs), 33,579 deletions, 31,198 insertions; 157 inversions and 86 duplications (Figure S1D-E). These numbers are in line with those reported from the three *M. m.* subspecies (*domesticus, musculus, and castaneus*)^34–36^, and support the assignment of the DDK strain to the *M. m. domesticus* lineage.

We focussed our analysis on the 476 kb *Om* critical interval which presents comparable variations to those observed genome-wide (Figure 1C and S1F). Interestingly, numerous non-synonymous mutations were detected in protein-coding genes, particularly in *Slfn* genes (Figure S1G), consistent with the extensive diversification previously reported in other mouse strains^24,35^. Specifically, the DDK genome displays multiple missense mutations in *Slfn1, Slfn3, Slfn4 and Slfn9*; a loss of *Slfn8* (see below); and a complete functional *Slfn10* gene (Figure 1C and S1G), despite its annotation as a pseudogene in the C57Bl/6 reference genome. Finally, *Slfn15*, which is misannotated as a long non-coding RNA (lncRNA) gene (*ENSMUSG00000121399*) in recent Gencode versions^37^ despite its high coding potential score and conserved open reading frame (ORF) in rodents^24^, carries a premature termination codon (PTC) in DDK, rendering it non-functional in this strain (Figure 1C and S1G).

We identified four deletions and three insertions larger than 1kb within the DDK *Om* region (Figure 1C, Supplementary Table S1). The largest deletion spans 35 kb between *Slfn1* and *Slfn4* and includes multiple direct repeats of highly homologous sequences. Another major deletion of 18.5 kb removes the entire *Slfn8* gene. Consistent with previous reports^34^, many of the SVs in the DDK *Om* region are associated with transposable elements. These include: i) the deletion of a full-length MERVL element upstream of *Slfn2*, also absent in the C3H/HeJ, 129S1, PWK/Ph and CAST/Ei inbred strains^35^; ii) tandem amplification of a pre-existing MERVL located downstream of *Slfn2* and upstream of its antisense lncRNA *Gm20234*; and 3) a DDK-specific MERVL insertion upstream of *Slfn1* that is absent in all other analyzed mouse strains (Figure 1C). To confirm the strain specificity of this insertion, we performed genomic PCR across a panel of mouse strains, including the DDK sister inbred strain DDD derived from the same dd outbred stock^38^. While the MERVL insertion was present in DDK, it was absent from all other strains tested, including DDD (Figure 1D and S2), establishing that this MERVL integrant is private to the DDK genome.

### The MERVL integrant drives *Slfn1* expression in DDK oocytes

To functionally dissect which of these genetic events underlie DKK incompatibility, we used the C.D-*Om^d^*^2^ (hereafter Omd2) congenic strain^39^, which carries a DDK-derived chromosome 11 fragment encompassing the *Om* locus on an otherwise incompatible BALB/c genetic background. Mouse Universal Genotyping Array (GigaMUGA)^40^ estimated the size of this DDK chromosomal fragment at 4 Mb (Figure S3A). Similar to DDK females, Omd2 females produce very few offsprings when mated with BALB/c males^39^ (see below), and the development of (Omd2xBALB/c)F1 embryos is compromised at the morula-to-blastocyst transition (Figure S3B). Live imaging revealed that these F1 embryos exhibit inefficient cavitation, characterized by repeated episodes of premature collapse of the blastocoelic cavity, ultimately followed by rapid degeneration of all embryonic cells (Figure S3C and Supplemental Movie S1).

Since the maternal *Om* factor is unique to DDK and deposited as an RNA in the oocyte cytoplasm^20^, we performed transcriptomic profiling of pooled f BALB/c or Omd2 metaphase II (MII) oocytes. Consistent with their genetic proximity, only 12 differentially expressed genes (DEGs; FDR<0.01 and |Log2(FC) |>2) were identified between Omd2 and BALB/c oocytes (Figure 1E, Supplemental Table S2). Two of these genes, *Slfn1* and *Gm20234,* are located within the *Om* critical region. *Gm20234* is a chimeric LTR-derived transcript antisense to *Slfn2* (Figure S4A). While its mis-regulation could reflect DDK-specific structural changes described above, its reduced expression in Omd2 oocytes argues against a role as the maternally deposited incompatibility factor. In contrast, *Slfn1* transcripts were strongly induced in Omd2 but undetectable in BALB/c oocytes, identifying *Slfn1* as a maternally expressed and deposited candidate from the DDK *Om* region (Figure S4B).

While DDK *Slfn1* transcripts were identical in DDK and BALB/c in spleen and thymus (Figure 1F), oocyte RNA-seq coverage extended 5’ of the canonical *Slfn1* transcription start site (TSS) into the MERVL FL copy in DDK (Figure 1F). MERVL elements are highly active during oogenesis and zygotic genome activation, and their LTRs have been widely co-opted as promoters in mammalian oocytes and early embryos^41^. We therefore hypothesized that the DDK specific MERVL insertion upstream of *Slfn1* may drive its expression in the oocyte. To test this hypothesis, we generated full-length cDNAs from Omd2 oocytes and used PCR amplification spanning *Slfn1* exonic sequence and a 5’ adapter to capture the oocyte-specific *Slfn1* 5’UTR (Figure S4C). Our results show that *Slfn1* transcripts in DDK oocytes initiate at the MT2-Mm LTR TSS and are spliced from the LTR splice donor site to two predominant cryptic splice acceptor sites located 162 and 303 nucleotides upstream of the canonical *Slfn1* TSS (Figure S4D).

Together, these findings demonstrate that the integration of a full-length MERVL element upstream of *Slfn1* in the DDK strain leads to its ectopic expression in oocytes.

### *Slfn1* expression in oocytes is responsible for DDK female incompatibility

To test whether *Slfn1* induction was required for incompatibility, we deleted the MERVL FL copy 5’ to *Slfn1* in the Omd2 strain using sgRNAs targeting unique flanking genomic regions, (Figure 2A and S5). The resulting Omd2-ΔMERVL strain displayed no detectable phenotypic abnormalities or reproductive defects. RNA-seq and quantitative RT-PCR analyses of individual MII oocytes from Omd2 or Omd2-ΔMERVL females revealed that *Slfn1* expression was abolished upon deletion of the MERVL copy (Figure 2B), unambiguously confirming that the MERVL integrant is solely responsible for driving oocyte *Slfn1* expression. We then mated Omd2 or Omd2-ΔMERVL females with BALB/c males and monitored offspring production over a 20-week period. Remarkably, whereas Omd2 females produced no viable offspring, Omd2-ΔMERVL females consistently produced normal litters (Figure 2C). Mean litter sizes of Omd2-ΔMERVL females were similar when mated with either compatible (Omd2-ΔMERVL) or incompatible (BALB/c) males (Figure 2D), indicating that *Slfn1* oocyte expression is responsible for the maternal incompatibility phenotype of the DDK syndrome.

**Figure 2:**
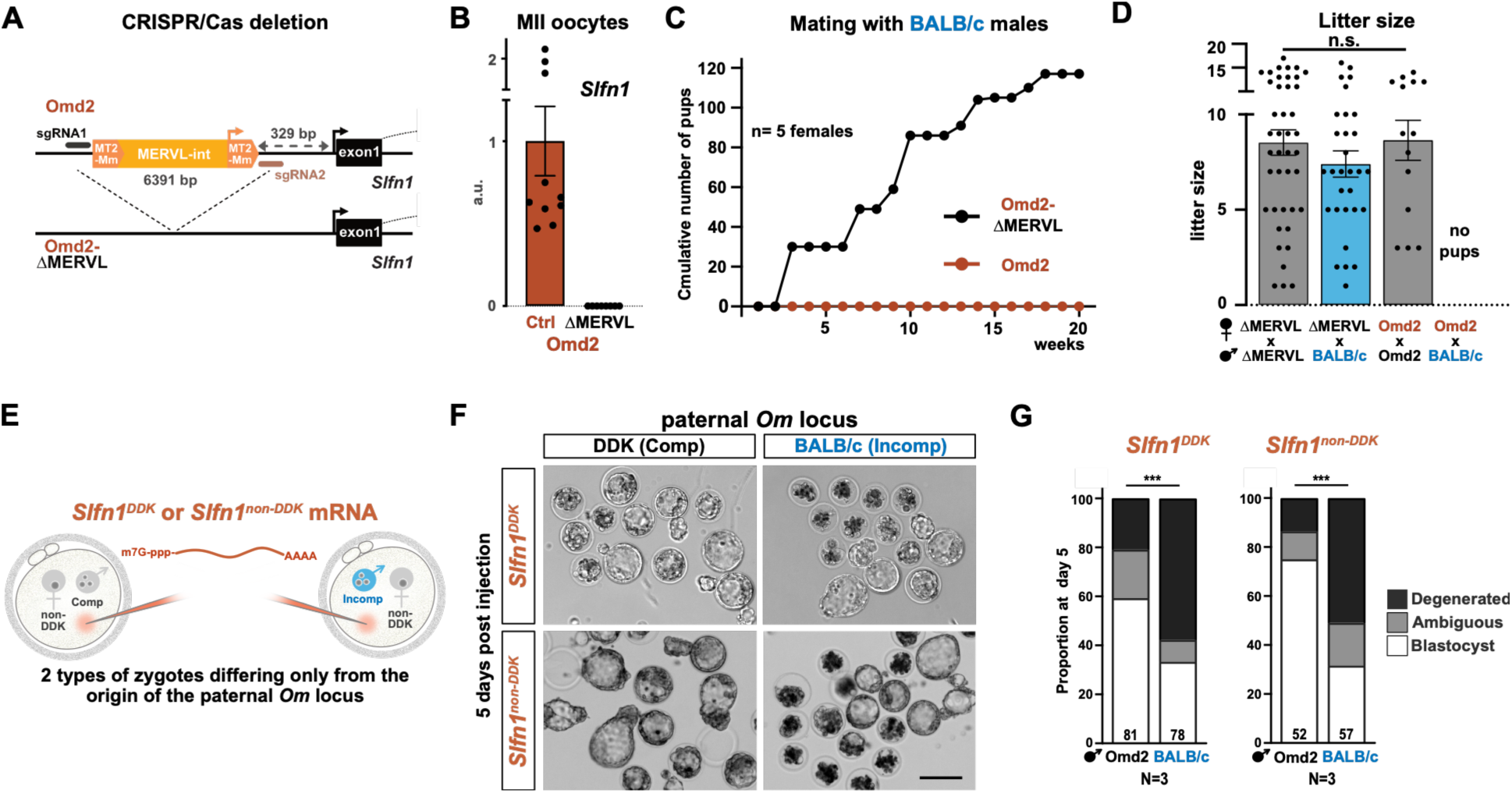
*Slfn1* expression in oocytes is both necessary and sufficient to cause hybrid incompatibility. **A.** Schematic representation of the CRISPR/Cas9 strategy used to the delete the full-length MERVL copy located upstream of *Slfn1* in the Omd2 strain. **B.** RT-qPCR quantification of *Slfn1* expression in single MII oocytes collected from Omd2 females carrying (Omd2-ΔMERVL) or lacking (Omd2, ctrl) the MERVL deletion. Mean level was arbitrarily set to 1 for Omd2. Error bars, SEM. **C.** Cumulative number of pups born over a 20-week period for Omd2 (n=5) and Omd2-ΔMERVL (n=5) females mated with BALB/c males. **D.** Litter sizes from the indicated crosses (5≤ n females ≤15). No pups were obtained when Omd2 females were mated with BALB/c males. Error bars, SEM. **E.** Scheme showing *in vitro* transcribed *Slfn1* mRNA microinjection into zygotes obtained from non-DDK (C57BL/6xDBA2)F1 females mated with either a compatible (Omd2) or incompatible (BALB/c) male. **F.** Brightfield images of microinjected embryos after 5 days in culture. Bar: 100µm. **G.** Developmental outcome of microinjected embryos. After 3 days in culture, embryos that had reached the morula were collected and their subsequent development monitored over an additional 2 days. Embryos undergoing blastocoelic cavity collapse without overt signs of cellular degeneration were classified as ambiguous, as they could represent either physiological transient collapse or early-stage degeneration. Differences in developmental outcomes between groups were assessed using a two-way chi-square test (p=1.2×10^-5^ and p=1.7×10^-5^, respectively). N: number of independent experiments.

To then determine whether ectopic *Slfn1* expression is sufficient to confer maternal incompatibility phenotype to a non-DDK background, we microinjected *in vitro* transcribed and polyadenylated *Slfn1^DDK^*mRNA into zygotes differing only in their paternal genome, originating from either incompatible (BALB/c) or compatible (Omd2) males (Figure 2E). After 72 hours in culture, similar proportions of both embryo groups reached the morula or early blastocyst stages (Supplementary Table S3). However, after an additional 48 hours, most embryos with a compatible paternal genome progressed to the blastocyst stage, while the majority of those with an incompatible paternal genome degenerated (Figure 2F, G and Supplementary Table S3). Together, these results demonstrate that *Slfn1^DDK^* mRNA is both necessary and sufficient to induce embryo lethality at the morula-to-blastocyst transition in incompatible crosses.

In previous work, we showed that *Slfn1* is subject to positive selection and represents the most divergent member of the *Slfn* family among Muroids^24^. Remarkably, the DDK *Slfn1* allele exhibits an unusual high number of non-synonymous substitutions (17 out of 337 amino acids; Figure S1G), far exceeding the divergence observed in more distantly related *Mus musculus* strains (*e.g. Mus musculus musculus* PWK: 0/337; *Mus musculus castaneus* CAST: 6/337). While the origin of this high sequence divergence remains unclear, it raises the possibility that SLFN1-driven incompatibility results from DDK-specific protein variant. To test this hypothesis, we repeated the microinjection assay using *Slfn1^C57Bl/^*^6^ mRNA and obtained the same results (Figure 2E-G and Supplementary Table S3), demonstrating that the SLFN1 incompatibility effect is not dependent on the DDK-specific protein sequence.

Altogether, our results demonstrate that *Slfn1* expression in oocytes is both necessary and sufficient to induce hybrid incompatibility.

### SLFN1 endonuclease activity causes translation inhibition and Integrated Stress Response activation in syndromic embryos

We first examined whether SLFN1 ribonuclease activity was required for (Omd2 x BALB/c)F1 embryos lethality. Structural studies have shown that, upon dimerization, the SCHLAFEN core domain forms an RNA-binding pocket with endonuclease activity, provided essential catalytic residues are present^42^. Among the five mouse SLFN12-related family members, SFLN1 harbours all residues predicted to support endonuclease activity, including the essential E193 catalytic residue required for RNA cleavage (Figure S6)^42–45^. Microinjection of *Slfn1^DDK^* mRNAs carrying an E193A mutation, *Slfn1^DDK-E193A^*, into zygotes with an incompatible paternal allele failed to induce developmental arrest (Figure 3A, B and Supplementary Table S4), demonstrating that an intact SLFN1 catalytic domain is required for embryonic toxicity.

**Figure 3:**
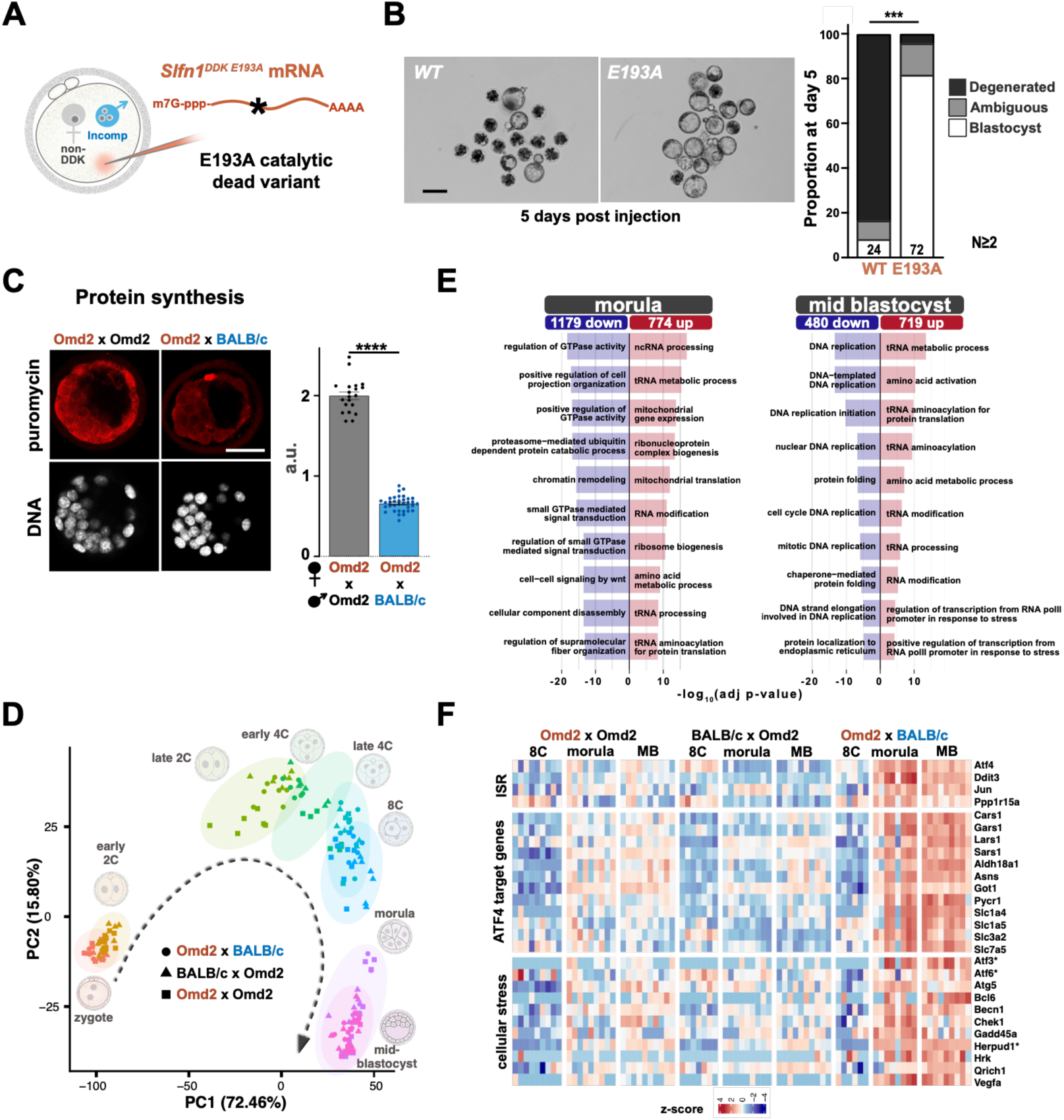
SLFN1 ribonuclease activity is required for translation inhibition and integrated stress response activation in syndromic embryos. **A.** Scheme showing *in vitro* transcribed *Slfn1^DDK-E193A^*mRNA microinjection. **B.** Brightfield images of embryos 5 days after microinjection of wild-type (WT) or catalytically inactive (E193A) *Slfn1* mRNA, together with their rates of development from morula to blastocyst stage. Differences in developmental outcomes were assessed using a two-way chi-square test (p=3.25×10^-14^). Bar: 100µm. N: number of independent experiments. **C.** Representative images and quantification of puromycin immunostaining (ribopuromycylation assay) in control Omd2 and (Omd2 x BALB/c)F1 syndromic embryos. Differences in puromycin incorporation levels were assessed using a two-sided Mann-Whitney test (p<0.001). **D.** Principal-component analysis of single-embryo RNAseq data from syndromic (Omd2 x BALB/c)F1 and non-syndromic control (Omd2 x Omd2 and (BALB/C x Omd2)F1) preimplantation embryos at multiple developmental stages. **E.** Top 10 enriched Gene Ontology biological process terms for upregulated and downregulated DEGs in syndromic embryos at the morula and mid-blastocyst stages. **F.** Heatmap showing a curated selection of DEGs associated with ISR, cellular stress pathways, or identified as direct transcriptional targets of ATF4. Genes marked with an asterisk are both associated with cellular stress and direct transcriptional targets of ATF4. Expression values were independently z-score normalized across samples for each gene.

Given the well documented role of SLFN proteins in tRNA binding and cleavage^25,29,31,32,46,47^, we next investigated whether protein synthesis was impaired in syndromic embryos. Using ribopuromycylation assays^48^ (Figure S7A), we found that syndromic embryos displayed markedly reduced levels of newly synthetized proteins compared to controls at early to mid-blastocyst stage (Figure 3C), indicating global translation inhibition. To test whether reduced protein synthesis might could account for developmental arrest ^49^, we inhibited translation in control embryos using puromycin before the morula-to-blastocyst transition and monitored their development by live imaging. Puromycin-treated embryos exhibited repeated episodes of blastocoelic cavity collapse closely recapitulating the phenotype of syndromic embryos (Figure S7B and Supplemental Movie S2).

To gain further mechanistic insight, we performed single-embryo transcriptomic profiling of (Omd2 x BALB/c)F1 syndromic embryos at multiple preimplantation stages: zygote, early and late 2-cell (E2C and L2C), early and late 4-cell (E4C and L4C), 8-cell (8C), morulae and mid-blastocyst (MB). Two control groups, embryos from BALB/c females mated with Omd2 males and embryos from Omd2 intra-strain crosses, were analysed in parallel. Principal component analysis of the individual embryo transcriptomes revealed that all three crosses follow similar preimplantation developmental trajectories, indicating that hybrid lethality in syndromic embryos is not caused by global defects in the early transcriptional waves governing early embryonic development (Figure 3D).

A distinctive transcriptional signature emerged in syndromic embryos at the morula stage preceding overt morphological defects (morula: 774 up, 1179 down; MB: 719 up, 480 down, Figure S8 and Supplementary Table S5). Gene ontology enrichment analysis revealed that downregulated genes were strongly enriched for pathways involved in small GTPase-mediated signal transduction, actin cytoskeleton organisation and Wnt signalling. By the blastocyst stage, this signature shifted towards pathways linked to DNA replication, and endoplasmic reticulum protein quality control (Figure 3E).

This temporal progression suggests that syndromic embryos are initially compromised in signalling and cytoskeletal processes essential for morphogenesis, followed by impaired cell proliferation and proteostasis. Conversely, genes involved tRNA processing and aminoacylation, amino acid metabolism and translation were upregulated at both stages (Figure 3C), potentially reflecting a compensatory response to impaired protein synthesis caused by tRNA cleavage. Consistent with previous reports showing that activation of SLFN11 or SLFN12 in human cells induces ribosome stalling, ribotoxic stress and integrated stress responses leading to apoptosis^32,33^, genes associated with the integrated stress response (ISR) and cellular stress pathways were also significantly upregulated in syndromic embryos at both morula and MB stages (Figure 3F). Notably, numerous transcriptional targets of ATF4, the main effector of the ISR, were some of the most strongly induced genes (Figure 3F).

Altogether, these findings establish SLFN1 ribonuclease activity as a key driver of hybrid incompatibility and implicate translation inhibition and ISR activation in the developmental failure of syndromic embryos.

### *Slfn15* is responsible for the paternal incompatibility

Previous studies have shown that the paternal factor must be expressed zygotically^14,17^. Indeed, transferring a DDK-incompatible pronucleus into an enucleated DDK zygotes prior to zygotic genome activation provokes the DDK syndrome. However, this effects only occurs when the non-DDK transferred pronucleus is of paternal, but not maternal, origin, suggesting that the gene encoding the paternal factor epigenetically acquires its “incompatibility potential” during spermatogenesis. Among the six genes in the *Om* region with significant expression in our RNA-seq dataset (mean TPM≥10 in at least one condition; Figure S9), *Slfn15* exhibited a particularly striking pattern, being first detected at the morula stage and significantly upregulated in syndromic morulae and mid-blastocysts compared to both control groups (Figure 4A). Allele-specific expression analysis revealed a strong bias toward the BALB/c allele in syndromic embryos indicating that most *Slfn15* expression originated from the paternal allele (Figure 4B and S10A). RT-qPCR of individual morulae and blastocysts further confirmed elevated *Slfn15* expression in syndromic embryos relative to controls (Figure 4C). Notably, BALB/c embryos displayed similarly high levels of *Slfn15* expression, consistent with the notion that increased *Slfn15* expression is attributable to the presence of a paternal BALB/c allele.

**Figure 4:**
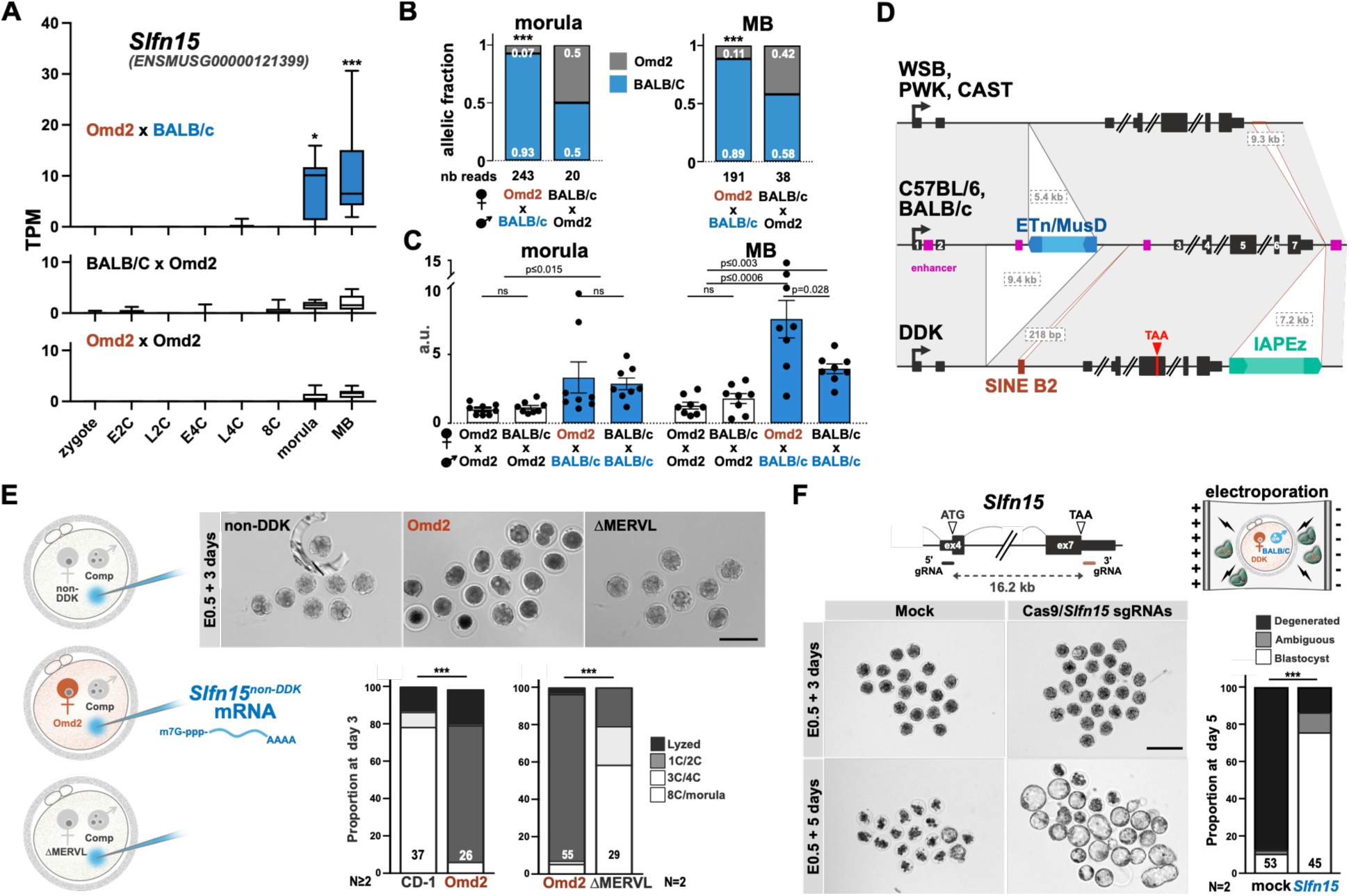
*Slfn15* is the paternal driver of the DDK syndrome. **A.** *Slfn15* mRNA levels at multiple developmental stages in embryos from the three crosses. Transcripts per million (TPM) values are presented as min-to-max box-and-whiskers plots. **B.** Allele-specific *Slfn15* expression analysis in (Omd2 x BALB/c)F1 and (BALB/c x Omd2)F1 embryos at the morula and mid-blastocyst stages. The number of informative reads from merged samples is indicated below each graph. **C.** RT-qPCR quantification of *Slfn15* expression in single morula and mid-blastocyst from Omd2, BALB/c and the two reciprocal F1 crosses. Mean expression level was arbitrarily set to 1 for Omd2 morulae. Differences in *Slfn15* expression levels were assessed using a two-sided Mann-Whitney test (p<0.01). Errors bars, SEM. **D.** Schematic representation of SVs present in *Slfn15* intron 2 and immediately downstream of the gene in various mouse strains. No SVs were identified between BALB/c and the mm39 C57BL/6 reference genome; both strains are incompatible with DDK. WSB, CAST and PWK are compatible strains. The predicted enhancers (filled pink boxes) from the Ensembl regulatory build are shown below the reference genome track. For clarity, a small 85bp deletion located immediately downstream of *Slfn15* in WSB, CAST and PWK has been omitted from the representation. **E.** Developmental outcome of zygotes microinjected with *in vitro* transcribed and polyadenylated *Slfn15* mRNA. Zygotes were obtained from non-DDK (BALB/c), Omd2 or Omd2-ΔMERVL females mated with a compatible (Omd2) male. Brightfield images and rates of development after 3 days in culture are also shown. Differences in developmental outcomes were assessed using a three-way chi-square test (p=2,18×10^-11^ and p=7.84×10^-9^, respectively). **F.** Schematic representation of the CRISPR/Cas9 electroporation strategy used to delete the *Slfn15* ORF in syndromic (Omd2 x BALB/c)F1 zygotes, using Cas9 and sgRNAs targeting genomic regions flanking the *Slfn15* ORF. Morula to blastocyst development rates of embryos electroporated with (*Slfn15* sgRNAs) or without (mock) sgRNAs are shown. Differences in developmental outcomes were assessed using a two-way chi-square test (p=7.51×10^-6^). Bar: 100µm. N: number of independent experiments.

Analysis of structural variants surrounding the *Slfn15* locus revealed that, compared with compatible CAST, PWK, and WSB strains, the incompatible BALB/c and C57BL/6 strains carry a 1.65 kb deletion immediately downstream of *Slfn15* and a 5.4 kb insertion within intron 2 of *Slfn15* (Figure 4D). Notably, the insertion corresponds to a full-length retrotransposon of the ETn/MusD family, known for its high transcriptional activity from the 8-cell stage onward^50,51^. Allele-specific expression analysis from published RNAseq datasets from embryos generated by hybrid crosses between C57BL/6 (with ETn/MusD) and CAST or JF1 (without ETn/MusD) ^52–54^ revealed that *Slfn15* transcripts are almost exclusively derived from the allele carrying the ETn/MusD insertion (Figure S10B,C). This suggests that functional *Slfn15* expression during preimplantation development may be absent in non-DDK compatible strains lacking this element. In the DDK strain, *Slfn15* has undergone multiple structural changes, including the downstream insertion of an IAPEz element, a 9.4 kb deletion encompassing regulatory elements and the ETn/MusD insertion, and the acquisition of a PTC (Figure 4D). Together, these observations suggest that hybrid incompatibility may depend on the production of a functional SLFN15 protein in early embryos and identify *Slfn15* as a compelling candidate for the paternal driver gene underlying the DDK syndrome.

To directly test this hypothesis, we produced *in vitro* transcribed and polyadenylated *Slfn15* mRNAs and microinjected them into zygotes differing only in their maternal genotype (Figure 4E). Three days after injection, only a small fraction of embryos with an Omd2 maternal genome advanced to the morula or early blastocyst stage; most arrested at the one- or two-cell stage or did not survive (Figure 4E). This early developmental arrest was not observed in zygotes with a non-DDK maternal genome (Figure 4E). To address whether this arrest results from incompatibility between maternally inherited *Slfn1* and zygotically provided *Slfn15,* we repeated the microinjection using zygotes derived from Omd2-ΔMERVL females, which lack oocyte *Slfn1* expression. In this context, *Slfn15* mRNA injection failed to induce developmental arrest (Figure 4E and Supplementary Table S6). Conversely, targeted deletion of *Slfn15* using Cas9-sgRNA electroporation in syndromic zygotes dramatically rescued embryonic development, with 29 out of 38 (76%) embryos reaching the blastocyst stage, and efficiently escaping the DDK syndrome (Figure 4F and Supplementary Table S7). Genotyping confirmed that 25 of26 (96%) blastocysts carried a deletion of the *Slfn15* ORF (Supplementary Table S8), indicating that loss of zygotic *Slfn15* expression is sufficient to restore normal preimplantation development.

Altogether, these results demonstrate that post-fertilization expression of *Slfn15* is both necessary and sufficient to induce preimplantation lethality maternally inherited *Slfn1*products are present.

### Co-expression of *Slfn1* and *Slfn15* is sufficient to induce ES cell death

To mechanistically investigate SLFN1-SLFN15 incompatibility, we established mouse embryonic stem (ES) cell lines expressing SLFN1-GFP constitutively, 3xFLAG-SLFN15 under doxycycline (dox) control, or both (Figure 5A). While expression of either SLFN1-GFP or 3xFLAG-SLFN15 alone had no notable impact on cell viability, their co-expression proved highly toxic, resulting in the death of most cells within 24 hours of dox induction (Figure 5B). Notably, this toxicity was completely abolished when catalytically inactive SLFN1^E193A^-GFP was co-expressed with 3xFLAG-SLFN15 recapitulating the catalytic requirement observed in embryos (Figure 5B). Early morphological signs of cellular stress and cell death emerged 6-8 hours after dox treatment (Figure S11A), coinciding with a marked decrease in global protein synthesis (Figure S11B)^28,52^. No translation inhibition occurred when the catalytically inactive SLFN1^E193A^ mutant was co-expressed with SLFN15 (Figure S11B). Although SLFN heterodimerization has not been previously reported, AlphaFold analysis predicted a high-confidence SLFN1-SLFN15 interaction (Figure S12), which we confirmed by co-immunoprecipitation (Figure 5C). Physical interaction was readily detected between SLFN15 and SLFN1^E193A^, and hardly with wild-type SLFN1, likely due to reduced SLFN15 protein synthesis in this context. Together these data suggest that SLFN1 and SLFN15 form heterodimers with ribonuclease activity inhibiting translation and thereby inducing cell death. We posit that these molecular functions underlie DDK zygotic hybrid incompatibilities.

**Figure 5:**
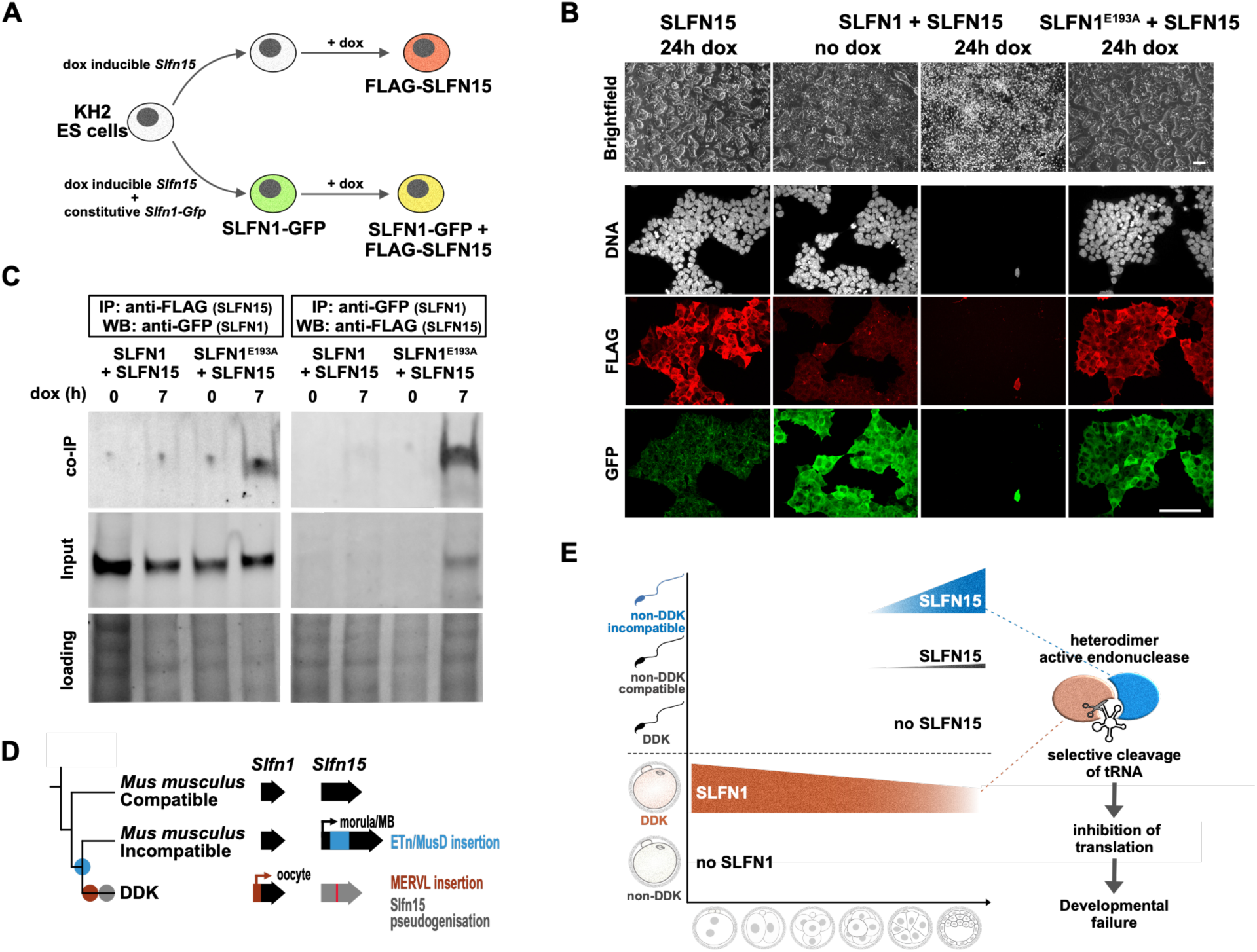
Co-expression of SLN1 and SLFN15 in ES cells induces rapid cell death. **A.** Diagram illustrating the strategy for obtaining ES cells expressing SLFN1 and SLFN15. **B.** Pictures of KH2 ES cells expressing 3xFLAG-SLFN15 under doxycycline (dox) control, SLFN1-GFP constitutively, or both. **C.** Co-immunoprecipitation of SLFN1-GFP and SLFN1E193A-GFP with 3×FLAG-SLFN15 in ES cells with or without dox treatment for 7 hours. SLFN1-GFP and SLFN1E193A-GFP were immunoprecipitated using an anti-GFP antibody, and 3×FLAG-SLFN15 was immunoprecipitated using an anti-FLAG antibody. **D.** Proposed evolutionary scenario for the origin of the DDK syndrome, illustrating the sequential retrotransposon insertion events that led to ectopic expression of *Slfn1* in oocytes and *Slfn15* in preimplantation embryos within the Mus musculus lineage. **E.** Model of the molecular mechanism underlying hybrid incompatibility in the DDK syndrome. Upon co-expression of maternally deposited SLFN1 and zygotically expressed SLFN15, heterodimerization leads to constitutive endoribonuclease activation, tRNA cleavage, global translation inhibition, integrated stress response activation, and ultimately embryonic death.

## DISCUSSION

Transposable elements are a major source of genomic novelty and have shaped key aspects of mammalian evolution, including brain development, placentation, and preimplantation embryogenesis^55–58^. Here, we describe a unique situation in which two independent and recent retrotransposon insertions rewire the expression of immune *Slfn* genes in the female germline and preimplantation embryo, ultimately causing postzygotic hybrid incompatibility between closely related mouse strains. This finding establishes retrotransposon-driven regulatory innovation at immune loci as a novel mechanism contributing to the emergence of reproductive barriers in mammals.

We provide a plausible evolutionary scenario for the origin of the DDK syndrome (Figure 5E). A first transposition event involving an ETn/MusD retrotransposon into intron 2 of *Slfn15* occurred within the *M. m. domesticus* lineage, leading to its ectopic expression in preimplantation embryos. Given that ETn/MusD elements are among the most highly expressed endogenous retroviruses in mouse preimplantation development from the 8-cell stage onward^51^, this insertion likely provides a regulatory element driving *Slfn15* expression during these stages. Although the precise timing of this insertion remains unknown, it must predate the human-mediated introduction of the house mice into the Americas, as DDK-incompatible (PERA) and DDK-compatible (PERC) inbred strains were derived from wild mice captured in Peru^18^. A second more recent event involved the integration of a MERVL element immediately upstream of *Slfn1*, which we show drives *de novo* oocyte expression of a chimeric *MERVL-Slfn1* transcript. This insertion likely occurred in the lineage leading to DDK, but its exact timing cannot be precisely determined. The linkage and phylogenetic distribution of both insertions support that the MERVL integrated into a chromosome already carrying the ETn/MusD insertion. We therefore propose that the homozygous fixation of this incompatible combination was facilitated by prior or subsequent suppression of *Slfn15* expression in DDK mice. Secondary suppression is supported by the accumulation of the multiple deleterious mutations we identified at the *Slfn15* locus, consistent with strong selective pressure to suppress SLFN15 function in preimplantation embryos. Another argument in favour of this hypothesis is the presence of a female meiotic drive on chromosome 11 that favours transmission of the DDK *Om* allele, which could have helped fixing the MERLV insertion despite its immediate fitness cost^59,60^. Nevertheless, whether these insertions became fixed because of an unknown beneficial effect of *Slfn* expression in oocytes, early embryos or elsewhere remains an open question.

Syndromic embryos resulting from crosses between DDK females and incompatible males are uniquely characterised by the simultaneous presence of maternally deposited SLFN1 and zygotically expressed SLFN15. The perdurance of maternal SLFN1 protein during preimplantation development, together with the onset of *Slfn15* expression, collectively dictates the timing of embryonic failure (Figure 5E). Our finding that premature expression of SLFN15, achieved through mRNA microinjection, results in earlier developmental failure indicates that the toxic interaction between SLFN1 and SLFN15 is rapidly manifested once both proteins are simultaneously present in the embryo. We demonstrate that the co-expression of the two proteins is sufficient to inhibit translation and trigger cell death, in a manner strictly dependent on SLFN1 catalytic activity. We postulate that, upon heterodimerization, SLFN1 and SLFN15 generate a constitutively active ribonuclease capable of cleaving tRNAs, thereby causing ribosome stalling, global translational repression, and ultimately cell death (Figure 5E). Although only homodimerization has been previously reported for SLFN family members, our demonstration of a functional SLFN1-SLFN15 heterodimer, together with our recent demonstration of ongoing diversifying selection at key residues within the SLFN dimerization interface in both rodents and primates^24^, suggests that the functional diversity and modes of action of SLFN proteins are broader than previously appreciated. Unlike longer SLFN members, whose ribonuclease activity requires activation by binding to nucleic acids or small molecules^32,61,62^, conformational change^46^ or dephosphorylation^63,64^, the endonuclease activity of the SLFN1-SLFN15 heterodimer appears to be constitutive. Whether heterodimerization with other family members similarly activates SLFN1 endonuclease activity, and how such activity is regulated in other biological contexts, particularly during immune responses and pathogen infection, remains important open questions.

Interestingly, prokaryotic Schlafen nucleases that target tRNAs during bacteriophage infection have recently been described^65,66^, suggesting the existence of an ancient tRNA-based defence mechanism conserved across the tree of life. Hundreds of prokaryotic proteins composed solely of a Slfn domain, analogous to mouse SLFN1, have been identified, yet their biological functions remain largely unknown.

## MATERIALS AND METHODS

### Mice

All experiments were performed in accordance with the French and European regulations on the care and protection of laboratory animals (EC Directive 86/609, French Law 2001-486 issued on June 6, 2001) and were approved by the Institut Pasteur ethics committee. All mice were maintained under a 14-hour light/10-hour dark cycle (lights on at 7:00 am). The C.D-*Om^d2^* congenic strain has been maintained in-house since its establishment by sibling matings, with occasional episodes of mating to BALB/cByJ animals followed by sibling matings to recover homozygosity at the *Om* locus. BALB/cByJ and (C57Bl/6 x DBA2)F1 mice were purchased from Janvier Labs and CD1 mice from Charles River Laboratories. The DDK strain has not been maintained as a live colony at the Institut Pasteur since 2012. Genomic DNA and RNA used in this study were therefore extracted from frozen DDK tissues.

### PCR on genomic DNA

Genomic DNAs from a collection of inbred strains representing the three *Mus musculus* subspecies were kindly provided by colleagues at the Institut Pasteur or purchased from the Jackson Laboratory (AKR/J, KK/HiJ, PERC/EiJ, RBA/Dn, Lewes/EiJ, PERA/EiJ, TIRANO/EiJ, and Zalende/EiJ). PCR amplification was performed using 20 ng of genomic DNA and the Taq Core Kit (MP Biomedicals, cat. 11EPTQK300) in a final reaction volume of 25 µl, containing 20.6 µl H₂O, 2.5 µl 10× buffer, 0.1 µl each of forward and reverse primers (100 µM), 0.5 µl dNTPs (10 mM each), and 0.2 µl Taq polymerase (5 U/µl). Thermal cycling conditions were as follows: initial denaturation at 95°C for 5 min, followed by 32 cycles of denaturation at 95°C for 10 sec, annealing at 60°C for 10 sec, and extension at 72°C for 1 min, with a final extension at 72°C for 10 min. PCR products were resolved on a 3% standard agarose gel (Invitrogen, cat. 16500-500) stained with GelRed (Interchim, cat. 41003) and electrophoresed in 0.5× ultrapure TAE buffer (Invitrogen, cat. 15558) using a RunOne electrophoresis unit (Embi Tec, cat. EP-2133).

### SNP Genotyping (GigaMUGA)

Genomic DNA was extracted from tail biopsies using Proteinase K (100 µg/ml; Eurobio, cat. GEXPRK01-B5) in lysis buffer containing 50 mM Tris-HCl pH 8.0, 100 mM NaCl, and 0.5% Tween-20, with overnight incubation at 56°C. Samples were then treated with RNase A (5 µg/ml; Roche, cat. 11119915001) at 37°C for 1 hour prior to phenol/chloroform extraction (Sigma, cat. P2069-100ML). DNA was precipitated by adding NaCl to a final concentration of 200 mM and 2.5 volumes of ethanol. After precipitation at -20°C for 24 hours, DNA pellets were resuspended in 50 µl of ultrapure DNase/RNase-free distilled water (Invitrogen, cat. 10977-035) by gentle pipetting using a cut pipette tip. DNA concentration was determined using a NanoDrop spectrophotometer, and integrity was assessed by agarose gel electrophoresis. DDK/Pas, BALB/cByJ and Omd2 DNA samples were shipped to Neogen Europe (Ayr, Scotland) on dry ice for genotyping via the GigaMUGA genotyping array (143,259 markers) on the Illumina Infinium platform(*67*). Analysis identified 29,233 and 191 polymorphic markers between DDK and BALB/c, and between Omd2 and BALB/c, respectively.

### Oocyte and embryo collection

MII oocytes were recovered from 4- to 5-week-old superovulated females by two successive intraperitoneal injections of 5 IU pregnant mare’s serum gonadotropin (PMSG; Centravet), followed 44–48 hours later by 5 IU human chorionic gonadotropin (hCG; Chorulon, Centravet). At 21–24 hours post-hCG injection, MII oocytes were collected from the oviducts and treated with hyaluronidase (Sigma, cat. H4272-30MG) to remove cumulus cells. After several washes in EmbryoMax M2 medium (Sigma, cat. MR-105D), oocytes from a single female were pooled, resuspended in a minimal volume of M2 medium and stored at −80°C until use.

Zygotes, late 4-cell, 8-cell, morula and mid-blastocyst stage embryos were obtained by natural matings of 6- to 12-week-old females, while early and late 2C as well as early 4C stage embryos were collected from 4- to 5-week-old superovulated females at 32–35 hours and 50–54 hours post-hCG injection, respectively. Correct developmental staging of early 2C, late 2C, early 4C and late 4C embryos was confirmed by the presence within the same litter of a minority of embryos at the immediately subsequent (late 2C and 4C) or preceding (early 2C and 4C) cell division stage.

### RNA in vitro synthesis and embryo microinjection

Microinjection experiments were performed as previously described in (*68*). The coding sequences for SLFN1^DDK^, SLFN1^C57BL/6^, SLFN1^DDK_E193A^, and SLFN15^C57BL/6^were synthesized by Proteogenix or Genscript and cloned together with *Globin* 5’ and 3’ UTRs(*69*) into pBluescript II SK(−) vectors. Plasmids were linearized with ScaI and used as templates for *in vitro* transcription using the mMESSAGE mMACHINE T7 Ultra Kit (Invitrogen, cat. AM1345). Capped and polyadenylated RNAs were purified by LiCl/ethanol precipitation and resuspended in Brinster’s buffer (10 mM Tris-HCl pH 7.5, 0.25 mM EDTA), and quality was assessed using an Agilent 2200 TapeStation system.

For microinjection, embryos were transferred to a depression slide in EmbryoMax FHM HEPES-Buffered Medium without Phenol Red (Sigma, cat. MR-025D) and microinjected using a Leica inverted microscope equipped with a Leitz micromanipulator. mRNA solutions at 500 ng/µl were injected into the cytoplasm of zygotes. Following microinjection, embryos were cultured in 10 µl drops of Embryo Love medium (Embryotech Laboratories, cat. ETECH-EL-20) covered with mineral oil (FertiPro, cat. MINOIL050) at 37°C 8% CO₂. After 3 days of culture, developmentally arrested embryos were scored and removed, and the progression of viable morulae or early blastocysts was monitored for an additional two days. Differences in the frequency of developmental progression (degenerated, ambiguous or blastocyst) were assessed using a two-sided Chi-square test.

### CRISPR/Cas9 zygote electroporation and embryo genotyping

Zygotes were electroporated using a NEPA21 electroporator fitted with a 5 µl CUY501P1-1.5 electrode chamber (Sonidel/Nepagene), loaded with Cas9/sgRNA ribonucleoprotein (RNP) complexes as described in (*70*) with some modifications. Briefly, stock solutions were prepared in Brinster’s buffer (10 mM Tris pH7.5; 0.25 mM EDTA pH8 filtered through a 0.45µm Minisart unit (Sartorius, cat. 16555-K)) at 150 pmol/µl and stored at −80°C. Immediately prior to use, each Alt-R CRISPR-Cas9 sgRNA (IDT) was diluted 1:25 (final concentration: 6 pmol/µl) and Alt-R S.p. Cas9 Nuclease V3 (IDT, cat. 1081058) was diluted 1:10 (final concentration: 6.2 pmol/µl) in Opti-MEM Reduced Serum Medium (51985-026, Gibco). For RNP complex assembly, 1 µl of the sgRNA (5’ and 3’) was diluted in 4 µl of Opti-MEM, and 1 µl of Cas9 was diluted in 9 µl of Opti-MEM. Each sgRNA-Cas9 complex was pre-assembled by combining 5 µl of diluted sgRNA with 5 µl of diluted Cas9 and incubating for 10 min at room temperature. The two RNP complexes were then mixed, and embryos were pre-incubated in a small drop of sgRNA/Cas9 solution. Ten to twenty zygotes were positioned at the center of the electrode chamber loaded with approximately 5 µl of the sgRNA/Cas9 solution, and sample impedance was measured and adjusted to 0.2–0.4 kΩ by adding or removing RNP solution as needed. Electroporation was performed using the following settings: poring pulse: 40 V, 3.5 ms length, 50 ms interval, 4 pulses, 10% decay rate, positive polarity; transfer pulse: 5 V, 50 ms length, 50 ms interval, 5 pulses, 40% decay rate, ±polarity, with the "current limit ON" option activated. Embryos were then recovered from the electrode chamber, rinsed several times and cultured at 37°C in 8% CO₂ in a non-treated 4-well plate (Thermoscientific, cat. 179820) containing 400 µl of Embryo Love medium per well. Electroporation of zygotes in Opti-MEM without RNP complexes served as a negative control and resulted in no significant developmental delay *in vitro*.

For MERVL deletion, 100 electroporated Omd2 zygotes were transferred into 5 pseudopregnant females giving rise to 18 newborns. PCR genotyping confirmed heterozygous or homozygous MERVL deletion in 13 out of the 17 pups genotyped. Two F0 females carrying the deletion were selected and crossed with Omd2 mice for two generations, followed by interbreeding to obtain Omd2-ΔMERVL animals homozygous for the MERVL deletion.

For *Slfn15* inactivation experiments, syndromic embryos that had not degenerated five days post electroporation were individually collected and genotyped using a single-embryo genotyping protocol described in(*71*). Briefly, each embryo was lysed in 15 µl of lysis buffer (10 mM Tris pH 8.0, 50 mM KCl, 0.01% gelatin, 300 µg/ml Proteinase K) at 56°C for 1 hour, followed by enzyme inactivation at 95°C for 10 min. Five µl were used for 21 cycles of pre-amplification using Phusion High-Fidelity DNA Polymerase (NEB, cat. M0530L) in a total volume of 25 µl (14 µl H₂O, 5 µl 5× buffer, 0.1 µl primers at 100 µM each, 0.5 µl dNTPs at 10 mM, and 0.20 µl Phusion polymerase). One µl of the pre-amplified product was then used for 21 cycles of nested PCR in a total volume of 25 µl using the same reagent concentrations. Both PCR steps used identical cycling conditions: initial denaturation at 98°C for 30 sec, followed by 21 cycles of 98°C for 15 sec, 60°C for 15 sec, and 72°C for 1 min, with a final extension at 72°C for 3 min.

### DDK genome sequencing

High molecular weight genomic DNA was extracted from DDK frozen tissues using the protocol described above for GigaMUGA genotyping except that DNA was recovered using a plastic rod rather than by centrifugation, to avoid mechanical shearing. Female genomic DNA was submitted to the Institut Curie genomic platform for library preparation and Illumina short-read whole genome sequencing. Oxford Nanopore Technologies (ONT) sequencing was performed at the Biomics platform of Institut Pasteur. Male genomic DNA was quantified using a Qubit fluorometer (Thermo Fisher Scientific), and quality-assessed using a Fragment Analyzer (Agilent Technologies).

For the first Nanopore sequencing run, DNA was size-selected by purification with 0.5x volume of AMPure XP magnetic beads (Beckman Coulter, USA) to remove fragments below 1kb. The library was prepared from 1 µg of purified genomic DNA using the Ligation sequencing DNA kit V14 (ONT, cat. SQK-LSK114) according to the manufacturer’s instructions. Library quality was assessed using a Qubit fluorometer and Fragment Analyzer. Sequencing was performed on a PromethION flow cell (FLO-PRO114M) using a P2 solo sequencer (ONT) for 72 hours. After 24h, the flow cell was washed and reloaded with fresh library using the Flow Cell Wash Kit (ONT, cat. EXP-WSH004) following the manufacturer’s instructions. Data were basecalled and demultiplexed with MinKNOW version 22.07.9 using Guppy version 6.3.9 and the super-accurate basecalling model. For the second sequencing run, the size-selection step was modified to enrich for longer reads. Briefly, 2 µg of genomic DNA were size-selected using the SRE XS kit (Pacific Biosciences) to remove fragments below 10kb, and libraries were prepared as described above. Basecalling and demultiplexing were performed using MinKNOW version 23.11.3 using Dorado version 7.2.12 and the High-accuracy model (400 bps). Mean read length was 1,620 bp for the first run and exceeded 12,000 bp for the second with N50 values of 9,150 bp to >23,000 bp respectively.

### Genome assembly, polishing and scaffolding

ONT basecalled reads were filtered using Filtlong (v0.2.1; --min_length 1000 --min_mean_q 9; https://github.com/rrwick/Filtlong). Illumina paired-end reads were adapter-trimmed using Trim Galore (v0.6.4; --paired --max_n 0; https://www.bioinformatics.babraham.ac.uk/projects/trim_galore/). For single nucleotide variant (SNV) calling, Illumina whole-genome sequencing data were processed with the Institut Curie VEGAN pipeline (v2.7.4; https://github.com/bioinfo-pf-curie/vegan; DOI: 10.5281/zenodo.20720451) running on Nextflow (v23.10.3). Reads were aligned to the mouse mm39 reference genome using BWA-MEM (v0.7.17)(72). Duplicate reads were marked with Picard (v3.0.0). Germline SNV and indel calling was performed with GATK HaplotypeCaller (v4.1.8.0)(*73*) with a minimum genotype quality of 30.

The DDK genome was assembled from filtered ONT reads using Flye(*74*) (v2.9.4-b1799; --nano-hq --genome-size 2.7g). The draft assembly was subsequently polished with Illumina short reads using POLCA(*75*, *76*) (v0.3.1; --careful). Assembly correction and scaffolding were performed using RagTag(*77*) (v2.1.0) in two successive rounds: first against the T2T C57BL/6J v1 reference genome(*78*) (GCA_964188535.1) for autosomes, and second against the T2T mhaESC v1.5 reference(*79*) (GCA_056825265.1) for sex chromosomes. Assembly quality and completeness were assessed using QUAST(*80*) (v5.3.0) and Merqury(*81*) (v1.3). Genome completeness was further evaluated based on the presence of conserved single-copy orthologs using BUSCO(*82*) (v6.0.0) with the mammalia_odb10 lineage dataset.

### Structural variant calling

ONT basecalled reads were aligned to the Mus musculus mm39 reference genome using Minimap2(*83*) (v2.30-r1287; -ax map-ont). Structural variants (SVs) were called from the aligned reads using three complementary long read SV callers: Sniffles2(*84*) (v2.7.2), SVIM(*85*) (v2.0.0), and cuteSV(*86*) (v2.1.3; --max_cluster_bias_INS 100 --diff_ratio_merging_INS 0.3 -- max_cluster_bias_DEL 100 --diff_ratio_merging_DEL 0.3). In parallel, assembly-based SV detection was performed by aligning the DDK genome assembly to mm39 using Minimap2 (v2.30-r1287; - ax asm5 --cs -r2k), followed by SV calling with SVIM-asm(*87*) (v1.0.3).

All SV callsets were filtered to retain only PASS-tagged variants using bcftools (v1.23). For read-base callers (Sniffles2, SVIM, and cuteSV), an additional minimum read support threshold of 3 was applied. The four filtered call sets were then merged using SURVIVOR(*88*) (v1.0.7; merge sample_files 1000 2 1 1 0 50), retaining only SVs supported by at least two callers and with a minimum length of 50 bp. The final merged call set was restricted to autosomes and sex chromosomes and limited to the following SV types: insertions (INS), deletions (DEL), inversions (INV), and duplications (DUP). SVs in the *Om* region were manually curated using local realignment to the reference genome using BLAST (https://blast.ncbi.nlm.nih.gov/)(89).

### Bulk RNA sequencing

Total RNA was extracted from spleen (DDK and BALB/c) and thymus (Omd2 and BALB/c) using the Precellys Lysis Kit CKMix 50-R (Bertin Technologies, cat. KT03961-1-013.2). Tissues were finely cut with scissors and transferred to a CKMix 50-R tube containing 1 ml of TRIzol (Thermo Fisher, cat. 15596026). Homogenization was performed at 4°C using the Precellys Evolution Plus equipped with the Cryolys cooling module, using the following program: 5,500 rpm for 20 sec, 30 sec pause, 5,500 rpm for 20 sec, 30 sec pause (2 ml tube setting). Samples were then incubated on ice for 2 min followed by 5 min at room temperature, and RNA extraction was carried out according to the TRIzol manufacturer’s instructions. Following isopropanol precipitation, RNA pellets were washed once with 70% ethanol and resuspended in ultrapure DNase/RNase-free water. RNA integrity and quality were assessed using an Agilent 2200 TapeStation and agarose gel electrophoresis. Poly(A)-selected RNA-seq libraries were prepared and sequenced as paired-end 150 bp reads on a NovaSeq6000 system (Illumina) by Novogene UK.

### RNA sequencing on pooled metaphase II (MII) oocytes

Total RNA was extracted from three biological replicates of pooled MII oocytes: 101, 130, and 107 Omd2 oocytes, and 156, 131, and 138 BALB/c oocytes, respectively, using the Arcturus PicoPure RNA Isolation Kit (Applied Biosystems, cat. 12204-01). ERCC RNA Spike-In control mix was added to each sample at a 1:10,000 dilution (Thermo Fisher/Invitrogen, cat. 4456740) prior to extraction but was not considered in subsequent computational analyses. RNA integrity and quality were assessed using an Agilent 2200 TapeStation. 5 ng of RNA were then processed by Novogene UK using the SMART-Seq™ v4 Ultra™ Low Input RNA Kit (Clontech). Briefly, first-strand cDNA was synthesized from total RNA and amplified by full-length LD-PCR. The resulting double-stranded cDNA was purified using AMPure XP beads and quantified with a Qubit fluorometer prior to sequencing library preparation. Sequencing was performed on an Illumina NovaSeq6000 platform using paired-end 150 bp reads.

### Single embryo RNA-seq

Single-embryo RNA extraction and library preparation were performed on individual mouse embryos as previously described in (*71*), based on the FLASH-seq method. Briefly, individual embryos were rinsed in PBS supplemented with 1 mg/ml acetylated BSA (Sigma-Aldrich, cat. B8894) at room temperature and transferred under a stereomicroscope to a PCR strip tube containing 5 µl of lysis buffer, minimizing BSA carryover. Amplification products were purified using 0.8x AMPure XP magnetic beads (Beckman Coulter, cat. A63881) and eluted in 15 μL of ultrapure DNase/RNase-free distilled water. Sample quality was assessed by qPCR using primer pairs for either TBP/Tcf25 or TBP/HPRT. Samples with a Ct value deviating by more than ±1 from the mean Ct value for a given developmental stage were excluded from further analysis. For each developmental stage and genotype, a minimum of five embryos from at least three independent litters were processed for library preparation using the Nextera XT DNA Library Preparation Kit (Illumina, cat. FC-131-1096). Sequencing was performed on Illumina NextSeq 2000 instruments across two runs, generating single-end reads of 70 bp (first run) and 80 bp (second run).

### Transcriptomic analyses

Paired-end FASTQ files (spleen, thymus and MII oocytes) were processed using a custom Julia-based trimming pipeline (SequenceTrimmer.jl and run_seq_trimmer.jl; https://github.com/TabithaRuecker/DDK.git) with the following parameters: reverse-complement alignment length set to 20 bp, maximum mismatch distance of 1, mismatch frequency threshold of 0.2, maximum of 5 ambiguous nucleotides, adapter seed length of 13 bp, and adapter consensus estimation performed on 100,000 reads. Adapter sequences were identified using the estimate mode, and reads were processed in blocks of 100 with no upper limit on the total number of reads analyzed. Trimmed paired-end reads were aligned to the GRCm39/mm39 reference genome using STAR (v2.7.11b). For standard transcriptomic analyses, a stringent alignment was performed, requiring unique genomic placement (--outFilterMultimapNmax 1) and allowing a maximum mismatch rate of 2.5% (--outFilterMismatchNoverLmax 0.025). As transposable elements and other repetitive sequences generate reads that frequently map to multiple genomic loci, a second, permissive alignment was generated for repetitive element analyses. This alignment allowed up to 100 mapping locations per read (--outFilterMultimapNmax 100; -- outSAMmultNmax 100) with relaxed mismatch filtering (--outFilterMismatchNoverLmax 0.05; -- outFilterMismatchNmax 4). For both alignments, splice junction annotation was incorporated through the STAR genome index (sjdbOverhang = 149), strand information was derived from intron motifs, and alignments were output as coordinate-sorted BAM files. Gene-level quantification was performed using featureCounts (v2.0.0) in unstranded mode (‘-s 0’), counting only reads mapping to exons (‘-t exon’) with a minimum mapping quality of 12 (‘-Q 12’), and using name as the feature identifier (‘-g gene_namè), and the Gencode annotation file gencode.vM36.annotation.gtf.

Differential gene expression analysis was performed using DESeq2(*90*) (v.1.48.2) on raw gene-level count data. A negative binomial generalized linear model (GLM) was fitted using the design formula ∼ Cross, comparing Omd2 and BALB/c oocytes. Statistical significance was assessed using the Wald test, and p-values were adjusted for multiple testing using the Benjamini–Hochberg method. Genes with an adjusted p-value (padj) < 0.01 and an absolute log2 fold change > 2 were considered differentially expressed. Positive log2 fold changes indicate higher expression in Omd2 oocytes, while negative log2 fold changes indicate higher expression in BALB/c oocytes.

For single-embryo sequencing reads, adapter trimming was performed using cutadapt (v. 4.9) to remove the Nextera adapter sequence (CTGTCTCTTATACACATCT) from the 3′ end of reads. Reads shorter than 25 bp after trimming were discarded (--minimum-length=25), bases with Phred quality scores below 20 were trimmed (--quality-cutoff=20) and reads containing more than 10% ambiguous nucleotides were removed (--max-n=0.1). Trimmed reads were then split into UMI and non-UMI reads using the custom Julia script split_fastq_UMInonUMI.jl (https://github.com/TabithaRuecker/DDK.git). Both read Both read types were aligned to the GRCm39/mm39 primary genome assembly using STAR (v2.7.11b) with a splice junction database overhang of 80 (--sjdbOverhang 80). Only uniquely mapped reads were retained (--outFilterMultimapNmax 1), allowing a maximum mismatch rate of 2.5% relative to read length (--outFilterMismatchNoverLmax 0.025). Gene-level read counting was performed using featureCounts (v2.0.0) with parameters -F GTF -Q 12 -t exon -g gene_name -s 0 and the GENCODE annotation file gencode.vM36.annotation.gtf.

Raw read counts were normalized by regularized logarithm (rlog) transformation using DESeq2 and subjected to principal component analysis (PCA). Differential gene expression analysis was performed separately for each developmental stage using the design formula ∼Cross. Differentially expressed genes (DEGs) were identified for each pairwise comparison using an adjusted p-value threshold of padj < 0.1 and classified as upregulated or downregulated based on the sign of the log2 fold change. Variance-stabilized expression values were obtained using DESeq2 and visualized as heatmaps using ComplexHeatmap, grouped by developmental stage (8-cell, morula, and mid-blastocyst) and cross type. A curated gene set associated with the integrated stress response (ISR)(*91*) or identified as direct target of ATF4(*92*) was selected for visualisation. Expression values were z-score normalized across samples for each gene prior to plotting. No hierarchical clustering was applied to rows or columns.

### Allele-specific expression analysis

Allele-specific expression analysis was performed on RNA-seq data from two types of hybrid preimplantation embryos: 1) our own dataset involving Omd2 and BALB/c strains and 2) published datasets involving C57BL/6 crossed with either CAST/EiJ(*52*, *53*) (GSE152103 and GSE278534) or JF1(*54*) (GSE259396) strains. Strain-specific SNP information was collected for DDK (this study), and for BALB/cByJ, CAST/EiJ and JF1/McJ(*36*). Only fixed, high-quality SNPs were retained (QUAL≥ 30 and INFO/DP≥ 10), yielding 2.5, 2.6, 21.3, and 21.1 million homozygous SNPs for DDK, BALB/cByJ, CAST/EiJ and JF1/McJ respectively. N-masked mm39 C57BL/6 reference genomes were generated using SNPsplit(*93*) (v. 0.6.0) with SNPs from either CAST/EiJ, JF1/MCJ or both DDK and BALB/cByJ SNPs.

SNP-aware alignment to the N-masked genomes was performed using STAR(*94*) (v. 2.7.11b) with dataset-specific parameters. For our dataset, STAR was run with the following parameters: -- outFilterMultimapNmax 1, --outFilterMismatchNoverLmax 0.025, --alignEndsType Local, -- outSAMmapqUnique 255, --outFilterScoreMinOverLread 0, --outFilterMatchNminOverLread 0, -- outFilterMatchNmin 0, --outSAMstrandField intronMotif, --outSAMattributes NH HI AS XS nM NM MD, to allow for uniquely mapped, stringent, local alignment of SNP-containing reads. For published datasets, parameters were adjusted to allow limited multimapping to retain reads from repetitive elements or paralogous regions, while allowing no mismatches: -- outFilterMultimapNmax 30, --outSAMmultNmax 30, --seedSearchStartLmax 20, -- outFilterMatchNmin 30, --outFilterMismatchNoverLmax 0, --alignEndsType EndToEnd, -- outSAMmapqUnique 255. Aligned reads were then assigned to their respective parental genomes using SNPsplit (v. 0.6.0).

Allele-specific read counts were quantified using featureCounts (subread v. 2.0.0), assigning primary alignments to genes based on the reference genome GTF annotation, without strand specificity and allowing reads overlapping multiple features. When comparing reciprocal crosses (own dataset and(*52*)), allelic imbalance was assessed using a two-sided Fisher’s exact test. For each gene, allelic read counts uniquely assignable to either parental genome were summed across all samples within each cross to construct a 2×2 contingency table of parental allele counts. The association between allelic ratio and cross direction was tested using a two-sided Fisher’s exact test (fisher.test, R stats package). Resulting p-values were adjusted for multiple testing across all genes using the Benjamini–Hochberg (BH) method. Genes were considered to show significant cross-dependent allelic bias at the following thresholds: * (padj < 0.05), ** (padj < 0.01), or *** (padj < 0.001). No additional effect-size threshold was applied for this comparison. In cases where only a single cross direction was available(*53*, *54*), allelic imbalance was assessed using a two-sided binomial test (binom.test, R stats package) against an expected 50:50 biallelic ratio, applied to pooled allelic read counts per gene and genotype. For GSE278534 (WT and matKO genotypes), male and female samples were pooled within each genotype prior to analysis; sex was not included as a covariate. For GSE259400, allelic imbalance was assessed independently at each developmental stage (morula, E3.5, and E4.5). P-values were adjusted for multiple testing across all genes within each genotype using the Benjamini–Hochberg method. To avoid classifying genes as allelically biased based solely on statistical significance driven by high sequencing depth, an effect-size threshold was applied in addition to the adjusted p-value criterion. Genes were classified as maternally or paternally biased only if the adjusted p-value was < 0.05 and the absolute deviation of the maternal (B6) allelic fraction from 0.5 exceeded 0.2 (i.e., maternal allelic fraction < 0.3 or > 0.7). Genes meeting the statistical significance criterion but not the effect-size threshold were classified as balanced or not significant. Significance levels are reported as * (padj < 0.05), ** (padj < 0.01), or *** (padj < 0.001), conditional on also meeting the effect-size threshold.

### Single oocyte/embryo RT-qPCR

cDNA from single oocytes and single embryos was obtained using the first step of the FLASH-seq protocol. Five microliters of 1:40-diluted cDNA were then used in duplicate for qPCR amplification with 4.75 µl of LightCycler 480 SYBR Green I Master Mix (Roche, cat. 04887352001) and 0.25 µl of forward and reverse primers (25 µM each), in a final reaction volume of 10 µl. Amplification was performed in 384-well LightCycler 480 plates (Sorenson Bioscience, cat. 38820) using a LightCycler 480 instrument (Roche). Thermal cycling conditions were as follows: initial denaturation at 95°C for 5 min (ramp rate 4.8°C/sec, 1 cycle), followed by 45 cycles of 95°C for 10 sec (ramp rate 4.8°C/sec), 61°C for 10 sec (ramp 2.5°C/sec), and 72°C for 25 sec (ramp rate 4.8°C/sec), with a final cooling step to 40°C for 1 sec (ramp 2.5°C/sec). Relative expression levels were determined using the Absolute Quantification/2nd Derivative Maximum method and normalized to the geometric mean of three reference genes (*TBP, Tcf25* and *HPRT*), selected for their stable expression across early preimplantation stages based on a published RNAseq dataset(*95*). All primers used were validated by standard curve analysis and melting curve generation to confirm amplification efficiency and the production of a single amplicon. Differences in gene expression were assessed using a two-tailed Mann-Whitney test.

### Ribopuromycylation

To assess active translation, elongating ribosomes were first stalled by incubating ES cells or embryos in medium supplemented with emetine (Sigma, cat. E2375) at 20 ng/µl for 10 min. Puromycin (Sigma, cat. P9620) was then added at 50 µg/ml for 5 min to label nascent polypeptide chains. Samples were subsequently fixed for 1 hour on ice with 4% paraformaldehyde (PFA) in 1x PBS alone (ES cells) or supplemented with 0.5% PVP (Sigma, cat. P0930-50G) and 0.5% Triton X-100 (Sigma, cat. T8787-250ML) (embryos). After permeabilization in 0.5% Triton X-100, 0.1% PVP and 1% Bovine Serum Albumin (BSA; Sigma, cat. A9418) in 1x PBS, puromycin incorporation was detected by immunostaining with a mouse monoclonal anti-puromycin antibody (BioLegend, cat. 381502) diluted 1:600 in staining buffer containing 0.1% Tween-20 (Sigma, cat. P1379) and 10% donkey serum (Sigma, cat. D9663) in 1x PBS for 72 hours at 4°C. Samples were then washed in a Nunclon Delta-treated 4-well plate (Thermo Scientific, cat. 176740) containing 400 µl PBS supplemented with 1% PVP and 0.1% Tween-20 per well for 30 min, and then incubated with a secondary DyLight 550-conjugated donkey anti-mouse secondary antibody (Jackson ImmunoResearch, cat. 712-475-151) and Hoechst 33342 (Thermo Fisher Scientific, cat. 62249) at a final dilution of 1:300. Puromycin incorporation levels were quantified by measuring the mean fluorescence intensity (MFI) for each sample. Differences in puromycin incorporation levels were assessed using a two-tailed Mann-Whitney test. To confirm that puromycin incorporation is dependent on active translation, a ribosome initiation inhibitor (harringtonine; Santa Cruz, cat. sc-204771) was applied at 2 µg/ml for 1.5 hours prior to emetine and puromycin treatment, which resulted in complete abolishment of the ribopuromycylation signal.

### Generation of ES cells lines expressing *Slfn1* and *Slfn15*

SLFN1^DDK^ and SLFN1^DDK_E193A^ ORFs were fused in-frame with GFP via a glycine linker (GSGGGGG) and cloned under the control of the CAG promoter, followed by an IRES-PuroR selection cassette. The resulting plasmids were introduced in KH2 ES cells(*96*) using Lipofectamine 2000 (Thermo Fisher Scientific, cat. 11668019). Six days post-transfection, GFP-positive cells were sorted by flow cytometry and plated at clonal density on mouse embryonic fibroblast feeder layers. Individual colonies displaying robust and homogeneous GFP expression were manually picked and expanded. The coding sequence for 3×FLAG-SLFN15^C57BL/6^ was synthesized by Genscript and cloned in pBS31 vector. KH2 ES cell lines constitutively expressing either SLFN1^DDK^-GFP and SLFN1^DDKE193A^-GFP were subsequently co-transfected with the pBS31-*3×FLAG-Slfn15^C57BL/6^* plasmid and a FLP recombinase expression plasmid, enabling site-specific integration of *3×FLAG-Slfn15^C57BL/6^* transgene at the *Col1a1* locus. Correct integration events were selected with hygromycin (50 µg/mL) for 10 days. Resistant colonies were expanded, and transgene inducibility was verified by doxycycline treatment followed by anti-FLAG immunofluorescence staining.

### ES Cell culture

Mouse embryonic stem cells (mESCs) were maintained on 0.1% gelatin-coated culture dishes in ES cell medium consisting of DMEM (Gibco, cat. 31966-021) supplemented with 15% fetal calf serum (FCS; Sigma, cat. F7524-500ML), 100 µM β-mercaptoethanol (Gibco, cat. 31350-010), and 10 ng/ml mouse leukemia inhibitory factor (LIF; Miltenyi Biotec, cat. 130-099-895). Cells were cultured at 37°C in a humidified atmosphere containing 5% CO₂. For routine passaging, cells were washed with PBS, dissociated with trypsin (Gibco, cat. 25300-054) for up to 5 min at 37°C, and neutralized with ES medium. Cell suspensions were centrifuged, resuspended in fresh medium, counted, and replated at the appropriate density on gelatin-coated dishes. For cryopreservation, cells were resuspended at 5× 10⁶ cells/mL in FCS supplemented with 10% DMSO (Sigma-Aldrich, cat. D2438-50ML), transferred to cryovials, frozen at -80°C, and subsequently stored in liquid nitrogen for long-term preservation. For thawing, frozen vials were rapidly warmed in a 37°C water bath, diluted in ES medium, centrifuged to remove DMSO, and replated on gelatin-coated dishes.

### Co-immunuprecipitation and western blotting

Two million cells were thawed from -80°C and directly lysed in a custom-made lysis buffer containing 10 mM Tris-HCl pH 7.5, 5 mM EDTA, 150 mM NaCl, 50 mM NaF (1 M stock; Sigma, cat. S7920), 10% glycerol, 1% NP-40 (Sigma, cat. 74385), and 30 mM sodium pyrophosphate. Immediately before use the buffer was supplemented with 1× Complete EDTA-free protease inhibitor cocktail (Roche, cat. 11873580001) and PhosSTOP phosphatase inhibitor (Roche, cat. 04906845001). Cell pellets (≤200,000 cells per10 µL of buffer) were lysed on ice in this buffer for two hours in the presence of benzonase (Merck, cat. 70664-3; 1:100–1:1000 dilution) to digest nucleic acids. Lysates were then sonicated for two cycles of 30 s on/30 s off using a Diagenode sonicator and centrifuged at 14,000 rpm for 4 min at 4°C. The clarified supernatant was collected and used for either co-immunoprecipitation or direct Western blotting.

For Western blotting, supernatants were quantified using the Qubit Protein Assay Kit (Thermo Fisher Scientific, cat. 10543343) and mixed with 4× Laemmli sample buffer (Bio-Rad, cat. 1610747). Approximately 20 µg of protein per sample was resolved on mPAGE 4–12% BisTris polyacrylamide gels (Millipore, cat. MP41G12) alongside a Precision Plus Protein Kaleidoscope prestained protein ladder (Bio-Rad, cat. 1610375). Proteins were transferred to nitrocellulose membranes using an Invitrogen iBlot 2 system (Invitrogen; NC Mini stacks, cat. IB23002). Transfer efficiency was verified by brief Ponceau S staining (Sigma, cat. P7170-1L), after which membranes were imaged and then washed once in PBS and blocked for two hours in 1× PBS containing 5% BSA. Primary and secondary antibody incubations were performed in 1× PBS containing 2.5% BSA. Primary antibodies were incubated for at least 16h at 4°C.

For co-immunoprecipitation using GFP-tagged proteins, samples were processed using ChromoTek GFP-Trap® Magnetic Agarose beads (cat. GTMAK) according to the manufacturer’s instructions. For co-immunoprecipitation using antibody-coupled beads, Protein G Sepharose Fast Flow beads (Sigma, cat. P3296-1ML) were first washed three times with TSE150 buffer (20 mM Tris-HCl pH 8.0, 150 mM NaCl, 2 mM EDTA, 1% Triton X-100, 0.1% SDS), with each wash consisting of a 5 min incubation at 4°C on a rotating wheel followed by centrifugation (1 min, 1,000 rpm). Beads were then blocked with BSA (Roche, cat. 10711454001) at 20 mg/ml for 4 hours at 4°C on a rotating wheel, washed three additional times with TSE150, and stored at 4°C as a 50% slurry in PBS until use.

For immunoprecipitation, 0.5 µL primary antibody was added to 500 µL of cell lysate and incubated overnight at 4°C on a rotating wheel. Twenty microliters of pre-blocked beads were then added and incubated for 2 hours at 4°C under rotation. Beads were collected by centrifugation (1 min, 1000 rpm), and the supernatant was retained as the flow-through fraction. Beads were washed once with 1 mL of lysis buffer for 5 min at 4°C on a rotating wheel, followed by four washes with TBS. Bound proteins were eluted by boiling the beads in 4x Laemmli sample buffer and centrifugation (10 min, 5000 rpm). Eluates, together with the corresponding input (5% of the initial lysate) and flow-through fractions were analysed by SDS–PAGE and immunoblotting.

The following primary antibodies were used for immunoprecipitation, immunoblotting, and immunofluorescence: mouse monoclonal anti-FLAG M2 (Sigma-Aldrich, cat. F3165), chicken polyclonal anti-GFP (Abcam, cat. ab13970), and mouse monoclonal anti-puromycin (BioLegend, cat. 381502). Primary antibodies were used at a 1:5,000 dilution for immunoblotting. The following secondary antibodies were used for immunofluorescence detection: Alexa Fluor 488-conjugated goat anti-chicken IgG (H+L) (Invitrogen, cat. A11039) and Alexa Fluor 488-conjugated donkey anti-mouse IgG (H+L) (Invitrogen, cat. A21202). Secondary antibodies were used at a 1:10,000 dilution for immunoblotting.

### Immunofluorescence staining

Cells cultured in µ-Slide 8-well chambers (ibidi, cat. 80826) were fixed with 4% paraformaldehyde (PFA) in 1x PBS for 1h on ice, washed three times with PBS, and stored in PBS until staining. Cells were permeabilised and blocked simultaneously in PBS containing 0.25% Triton X-100 and 1% bovine serum albumin (BSA) for 30 min at room temperature. After washing with PBS containing 0.1% Tween-20 (PBS-T), samples were incubated for two days with primary antibodies diluted in PBS-T. Samples were then washed three times with PBS-T and incubated for 2h at room temperature with the appropriate secondary antibodies, donkey anti-mouse Alexa Fluor 488 (Invitrogen, cat. A21202) and goat anti-chicken Alexa Fluor 488 (Invitrogen, cat. A11039), together with Hoechst 33342 (1:1,000) diluted in PBS-T. Following secondary antibody incubation, samples were washed thoroughly with PBS-T and imaged directly in the chamber slides.

### Confocal microscopy and live imaging

All images were acquired using a Zeiss LSM 900 confocal microscope. Embryos were placed in an in-house-designed and -fabricated eggbox imaging device(*68*) to facilitate multi-position imaging and scanned using a Plan-Apochromat 20x air objective. Images were acquired using bi-directional scanning with 2x zoom and a 1 Airy unit pinhole at the optimal confocal resolution. Z-stacks were acquired with a 2 µm step size, generating 16-bit images (1,293 x 1,293 px). For live imaging, embryos were maintained in a humidified incubation chamber at 37°C and 8% CO₂ and imaged at 10-minute intervals over a total period of 24 to 48 hours.

### Structural alignment of mouse SCHLFAFEN proteins

Coding sequences of mouse *Schlafen* RefSeq mRNAs were translated into protein sequences using the following accession numbers: *mSlfn1* (NM_011407.2), *mSlfn2* (NM_011408.1), *mSlfn3* (NM_011409.1), *mSlfn4* (NM_001302559.1), *mSlfn5* (XM_017314615.1), *mSlfn8* (NM_181545.4), *mSlfn9* (XM_006533235), *mSlfn10* (NR_073523.1), *mSlfn14* (NM_001166028.1), and *mSlfn15* (manually assembled). The resulting protein sequences were then aligned to human SLFN11 (translated from RefSeq mRNAs NM_152270.4) used as a reference, using MAFFT (v1.5.0)(*97*).

### Counting non-synonymous mutations at coding genes in the *Om* locus

Exonic DNA sequences were extracted from the longest annotated isoform of each coding gene in the *Om* locus (*Slfn1*: NM_011407.2, *Slfn2*: NM_011408.1, *Slfn3*: NM_011409.1, *Slfn4*: NM_001302559.1, *Slfn5*: XM_017314615.3, *Slfn8*: NM_181545.4, *Slfn9*: XM_006533235.5, *Slfn10*: NR_073523.1, *Slfn14*: NM_001166028.1, *Slfn15*: manually assembled, *Ap2b1*: NM_001035854, *Pex12*: XM_030245452.1, *Rasl10b*: NM_001013386) and mapped onto the C57BL/6 *Om* locus to determine their genomic coordinates. Exon annotations were then manually created in Geneious Prime based on these coordinates. The *Om* genomic region was extracted from the BALB/cByJ, 129S1/SvImJ, WSB/EiJ, PWK/PhJ, CAST/EiJ, and DDK genome assemblies(*34*) and aligned to the C57BL/6J reference genome using MAFFT v1.5.0. Exon annotations were subsequently transferred across strains using the Transfer Annotations function in Geneious Prime (v2026.0.2). Coding sequences for each strain were manually reconstructed based on the transferred exon annotations, translated into protein sequences, and aligned to quantify amino acid substitutions relative to the C57BL/6 reference.

### Structure Prediction and Alignment

Structural alignment between human SLFN11 (PDB: 7ZEL) and AlphaFold3-predicted mouse SLFN1 was performed using the Matchmaker tool implemented in UCSF ChimeraX (v1.10.1). The interaction between C57BL/6 SLFN15 and DDK SLFN1 was predicted using AlphaFold(*98*). Residues were considered to be in contact if they were located within 4 Å of each other and if the predicted aligned error (PAE) was below 4 Å. PAE matrices were visualized by uploading the structure (.cif) and scores (.json) output files from AlphaFold3 into the PAE Viewer web tool (https://pae-viewer.uni-goettingen.de/).

### Statistical analysis

All statistical analyses were performed with Graphpad prism 11.0 software or R Bioconductor. The tests used are indicated in individual method paragraphs as well as in figure legends. *p< 0.05,**p< 0.01, ***p< 0.001, ****p< 0.0001.

## Acknowledgments

We are grateful to the staff of the animal facility of Institut Pasteur for animal care and their help during this work, Chloé Baum and Elodie Turc from the Biomics Platform (C2RT, Institut Pasteur, supported by France Génomique (ANR-10-INBS-09) and IBISA) for Nanopore sequencing, and Emmanuel Frachon and Samy Gobaa from the Biomaterials and Microfluidics platform (C2RT, Institut Pasteur) for the production of eggbox imaging devices. We also thank Xavier Montagutelli for critical reading of the manuscript, Sabrina Coqueran for her help in collecting MII oocytes, Eskeatnaf Mulugeta for his help in assembling llumina short reads, Xavier Montagutelli for his help in the analysis of GigaMUGA data, Jean-Louis Guénet, Franck Bourgade, Xavier Montagutelli, Caroline Manet, Laurine Conquet and Jean-Jaubert for providing genomic DNAs from various mouse strains. This article is dedicated to the memory of Charles Babinet and Jean-Louis Guénet.

## Authors contributions

T.R., S.V.-P., Conception and design, Acquisition of data, Analysis and interpretation of data; J.M., L.L., A.T., Analysis and interpretation of data. P.N., Analysis and interpretation of data, Drafting or revising the article; D.B., A.M., M.C.-T., Conception and design, Analysis and interpretation of data, Drafting or revising the article.

## Data and code availability

The DDK genome sequencing data have been deposited at DDBJ/ENA/GenBank database under BioProject number PRJNA1492211. All RNA-seq data generated in this study are available in the Gene Expression Omnibus database under accession numbers GSE338323. Published RNA-seq data were obtained from the GEO series GSE152103, GSE278534 and GSE259396. All softwares used for analysis are freely available and can be found in the methods section. For RNAseq analysis, code can be found on: https://github.com/TabithaRuecker/DDK.git. All other relevant and details of resources can be found in the main text or the supplementary materials.

---

**Figure S1:**
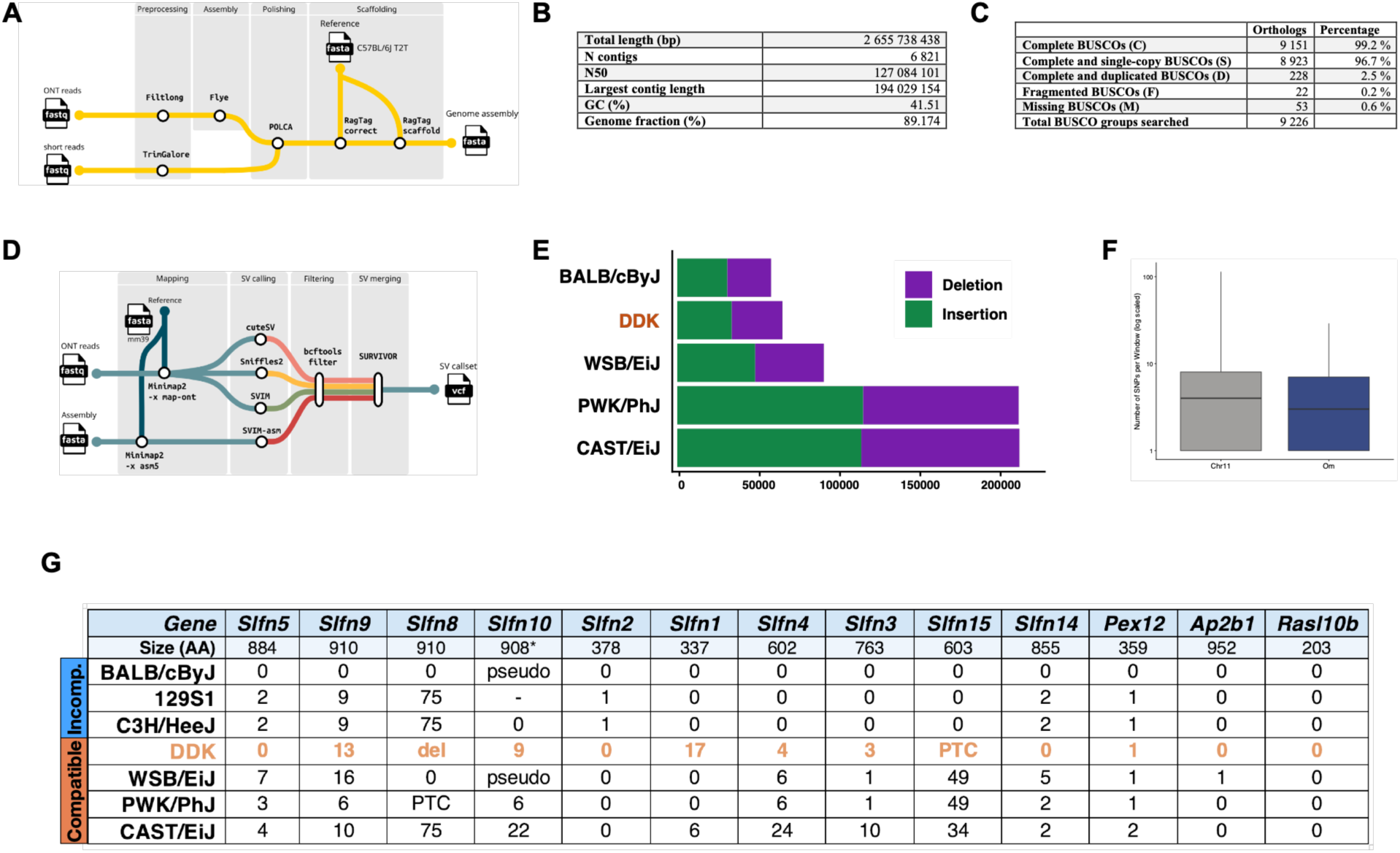
*De novo* DDK genome assembly reveals extensive structural variation and numerous non-synonymous variants in *Slfn* genes. **A.** Workflow of the DDK genome and annotation pipeline, including ONT long-read and Illumina short-read sequencing, polishing, and scaffolding steps. **B.** QUAST assembly quality statistics for the DDK genome assembly evaluated against the T2T C57BL/6J reference genome. **C.** BUSCO completeness analysis of the assembled DDK genome. **D.** Workflow of the structural variant (SV) calling pipeline, combining three long-read SV callers and assembly-based SV detection. **E.** Total number of SVs identified in DDK relative to the *Mus musculus* GRCm39 (C57BL/6J) reference genome, compared to the numbers of SVs identified in a selection of other inbred strains. **F.** DDK SNP density within 1 kb windows across chromosome 11 and within the *Om* critical region. **G.** Non-synonymous mutations identified in the ORF of protein-coding genes within the *Om* region, relative to C57BL/6J (and to 129S1 for *Slfn10*, which is annotated as a pseudogene in the reference genome). For each gene, the number of amino acid substitutions and the presence of a premature termination codon (PTC), a deletion (del) or pseudogenization (pseudo) are indicated, based on the DDK genome assembly and available genome sequences from a selection of compatible and incompatible strains.

**Figure S2:**
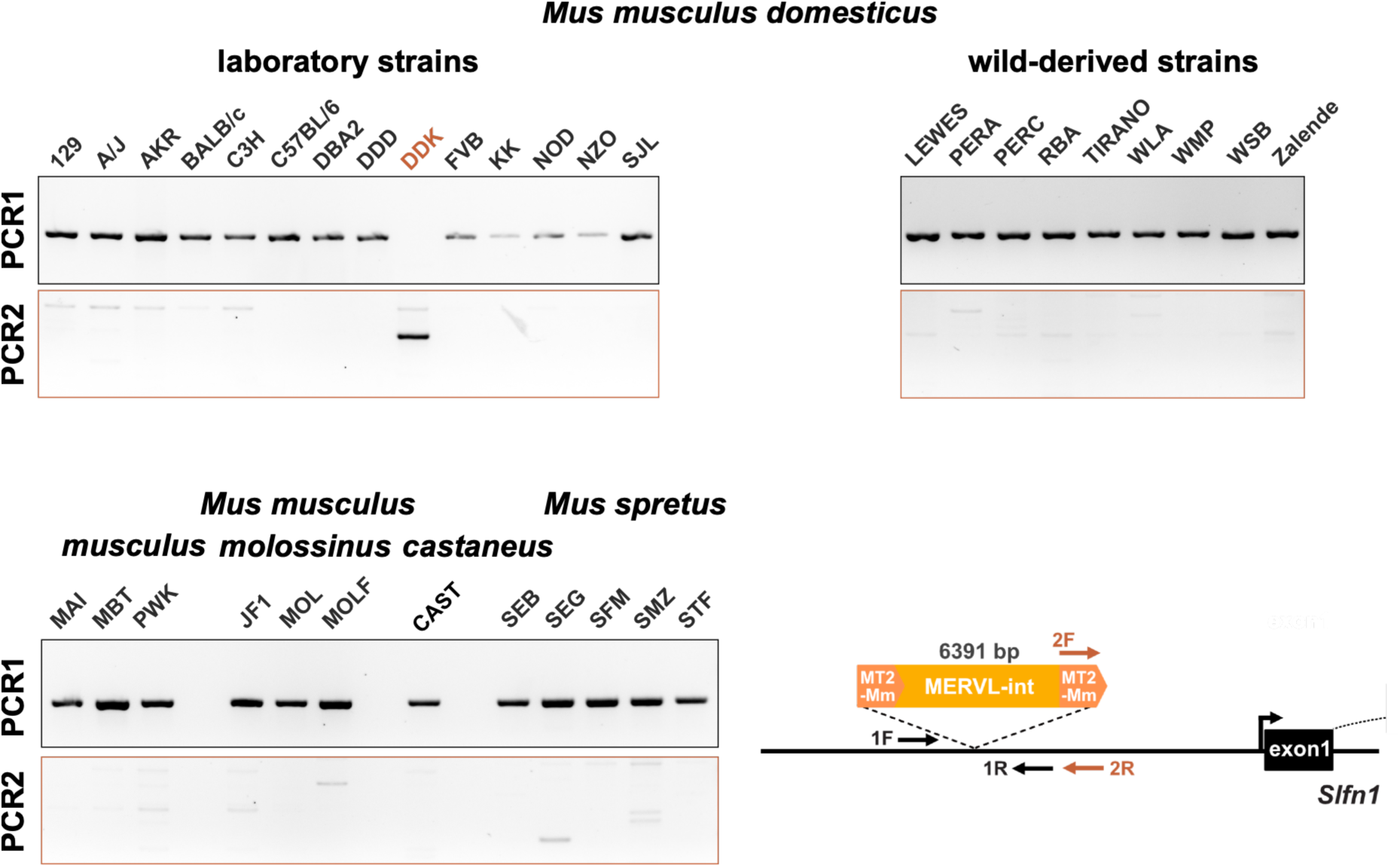
The full-length MERVL insertion upstream of *Slfn1* is unique to the DDK strain. Genomic PCR was performed on a panel of inbred mouse strains using primers flanking the MERVL integration site (PCR1) or primers specific to the MERVL insertion (PCR2).

**Figure S3:**
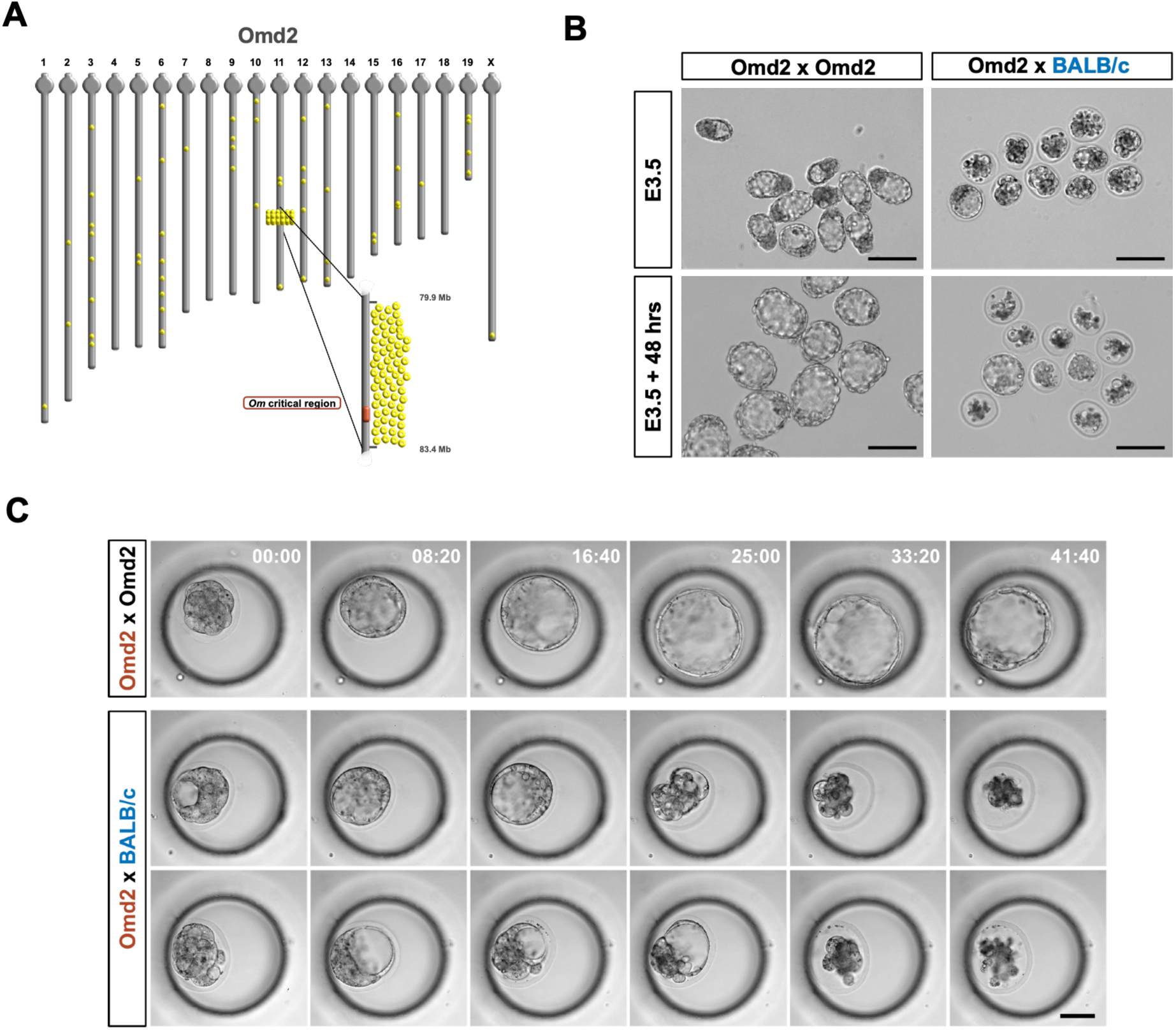
The Omd2 congenic strain faithfully recapitulates the DDK incompatibility phenotype. **A.** Genome-wide distribution of GigaMUGA genotyping array probes carrying a DDK-specific SNP (yellow dots) in the Omd2 congenic strain. A continuous block of approximately 4 Mb of DDK haplotype (70 out of 162 non-ambiguous SNPs of DDK origin) is identified at the distal region of chromosome 11, encompassing the *Om* critical region. The remaining DDK-derived SNPs are isolated and broadly distributed across all chromosomes, reflecting the BALB/c genetic background. **B.** Brightfield images of embryos collected at E2.5 from syndromic (Omd2 x BALB/c) or (Omd2 x Omd2) control crosses and cultured *in vitro* for 2 days. Most syndromic embryos degenerated, while most control embryos developed to the blastocyst stage and hatched from the zona pellucida. **C.** Selected frames from Supplemental Movie S1 illustrating the initial expansion of the blastocoelic cavity followed by its rapid collapse and the progressive degeneration of blastomeres in the syndromic embryos. Bar: 100µm.

**Figure S4:**
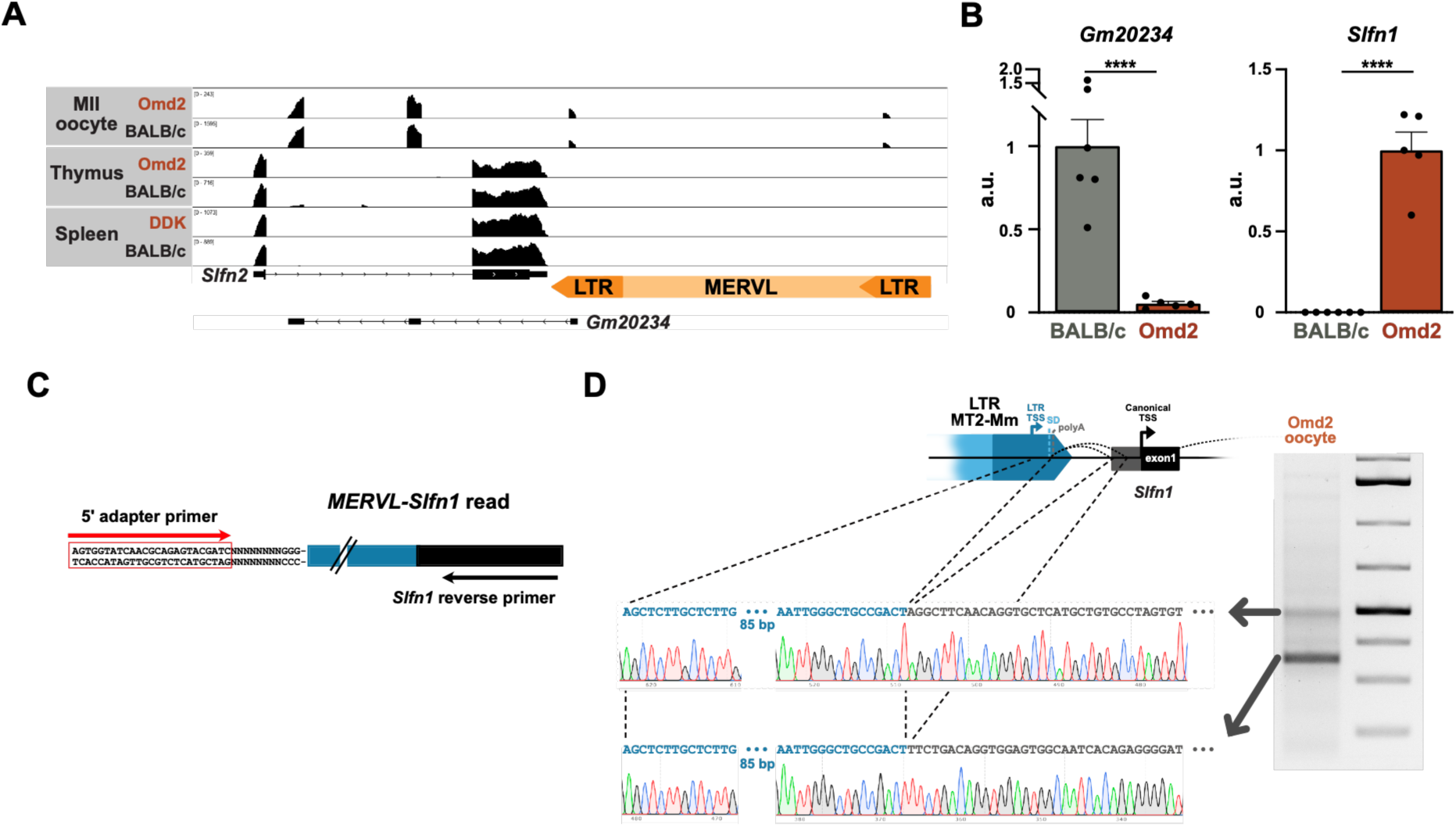
Structure and expression of *Gm20234* and *Slfn1* transcripts in DDK and BALB/c oocytes. **A.** IGV plots showing *Gm20234* and *Slfn1* read coverage after alignment to the GRCm39 reference genome. Reads mapping to *Gm20234* exon1 are distributed across the two identical LTRs of the full-length MERVL element located upstream. **B.** RT-qPCR quantification of *Gm20234* and *Slfn1* expression in single MII oocyte collected from BALB/c and Omd2 females. Mean expression levels of *Gm20234* and *Slfn1* were arbitrarily set to 1 for BALB/c and Omd2 respectively. Differences in expression levels were assessed using a two-sided Mann-Whitney test (p<0.0001). Error bars, SEM. **C.** Schematic representation of the 5’ extremity of the *MERVL-Slfn1* chimeric cDNA generated by the Flash-seq protocol. D. Sanger sequencing results of the two main PCR products obtained by amplification of Omd2 oocyte Flash-seq cDNAs using a 5’ adapter primer and a *Slfn1*-specific reverse primer. The two splicing events between the MT2-Mm LTR splice donor and cryptic splice acceptor sites located upstream of the canonical *Slfn1* TSS are illustrated on the schematic. SD and polyA: splice donor and polyadenylation signal present in the MT2-Mm LTR.

**Figure S5:**
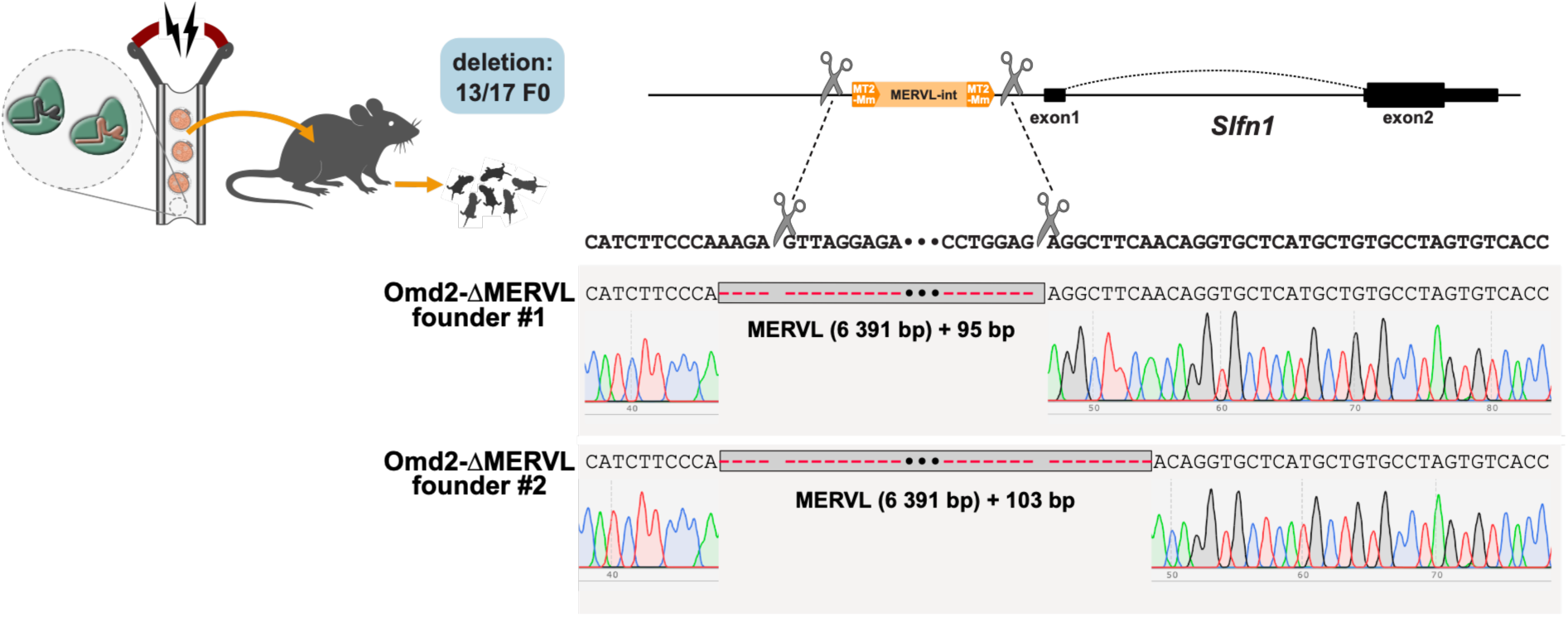
Generation of Omd2-ΔMERVL founder mice by CRISPR/Cas9 electroporation of Omd2 zygotes. Two founder mice carrying the targeted MERVL deletion were selected from 13 edited newborns. Sanger sequencing of the targeted region revealed that the deletion in founder #1 spans the genomic interval between the two CRISPR/Cas9 cut sites, with an additional 4 bp deletion at the 5’ end. The deletion in founder #2 encompasses that of founder # 1 and extends an additional 8 bp at the 3’ end. Both deletions result in complete removal of the full-length MERVL copy located upstream of *Slfn1*.

**Figure S6:**
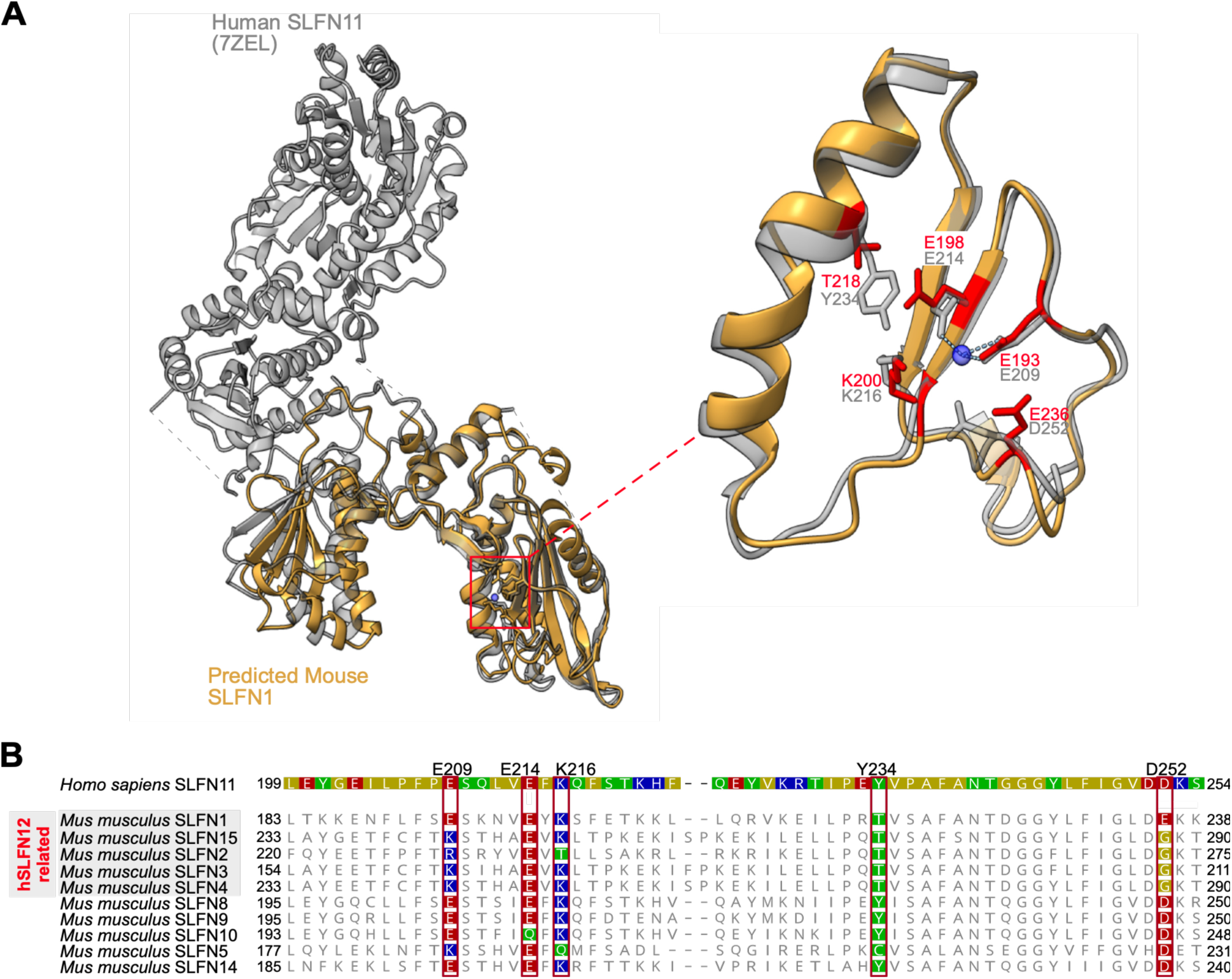
Identification of SLFN1 catalytic residues by structural and sequence analysis. **A.** Superposition of the cryo-EM structure of human SLFN11 (PDB: 7ZEL) with the AlphaFold3-predicted structure of mouse SLFN1. A close-up view of the nuclease active site is shown on the right, highlighting the key catalytic residues of hSLFN11 (E209, E214, K216, Y234 and D252) and their predicted structural equivalents in mSLFN1 (E193, E198, K200, T218, and E236). **B.** Multiple sequence alignment of mouse SLFN family members with hSLFN11 in the region surrounding the nuclease active site. Among the short, hSLFN12-related mouse SLFN members, SLFN1 is the only one retaining the critical glutamic residue (E209 in hSLFN11), while all other short members lack this residue and are predicted to be catalytically inactive. Among long mouse SLFN members, all except SLFN5 are predicted to retain ribonuclease activity based on conservation of the catalytic residues.

**Figure S7:**
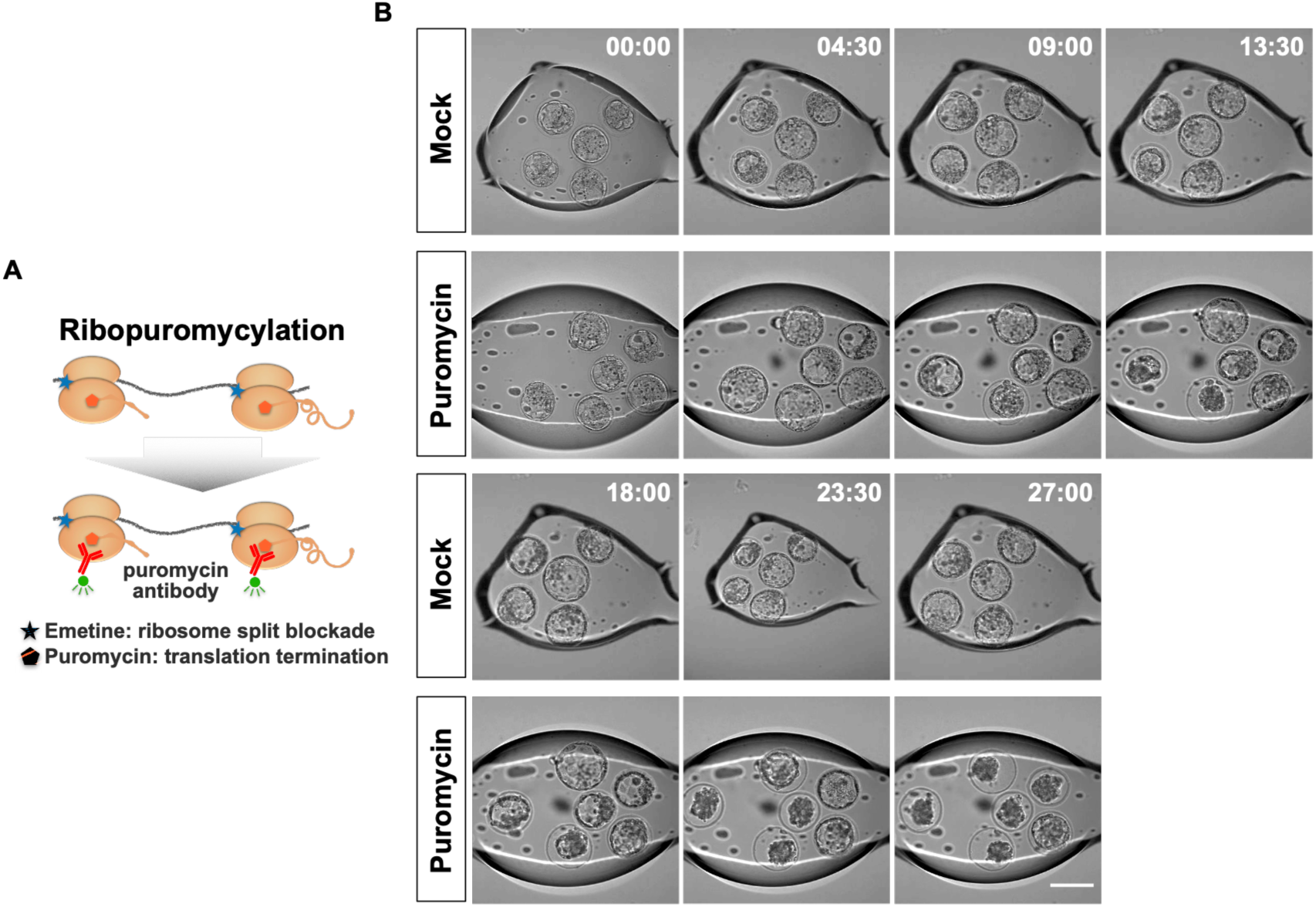
Quantification and experimental inhibition of protein synthesis in preimplantation embryos. **A.** Schematic representation of the ribopuromycylation assay. Following treatment with emetine, which prevents ribosome translocation and dissociation of nascent polypeptides, puromycin is added to label nascent peptides trapped on the ribosomes. Puromycin-labeled nascent peptides are then detected by immunostaining with an anti-puromycin antibody, providing a quantitative readout of active translation. **B.** Selected frames from Supplemental Movie S2 illustrating the morula to blastocyst development of control CD1 embryos cultured in the presence or absence (mock) of puromycin (2µg/ml). Bar: 100µm.

**Figure S8:**
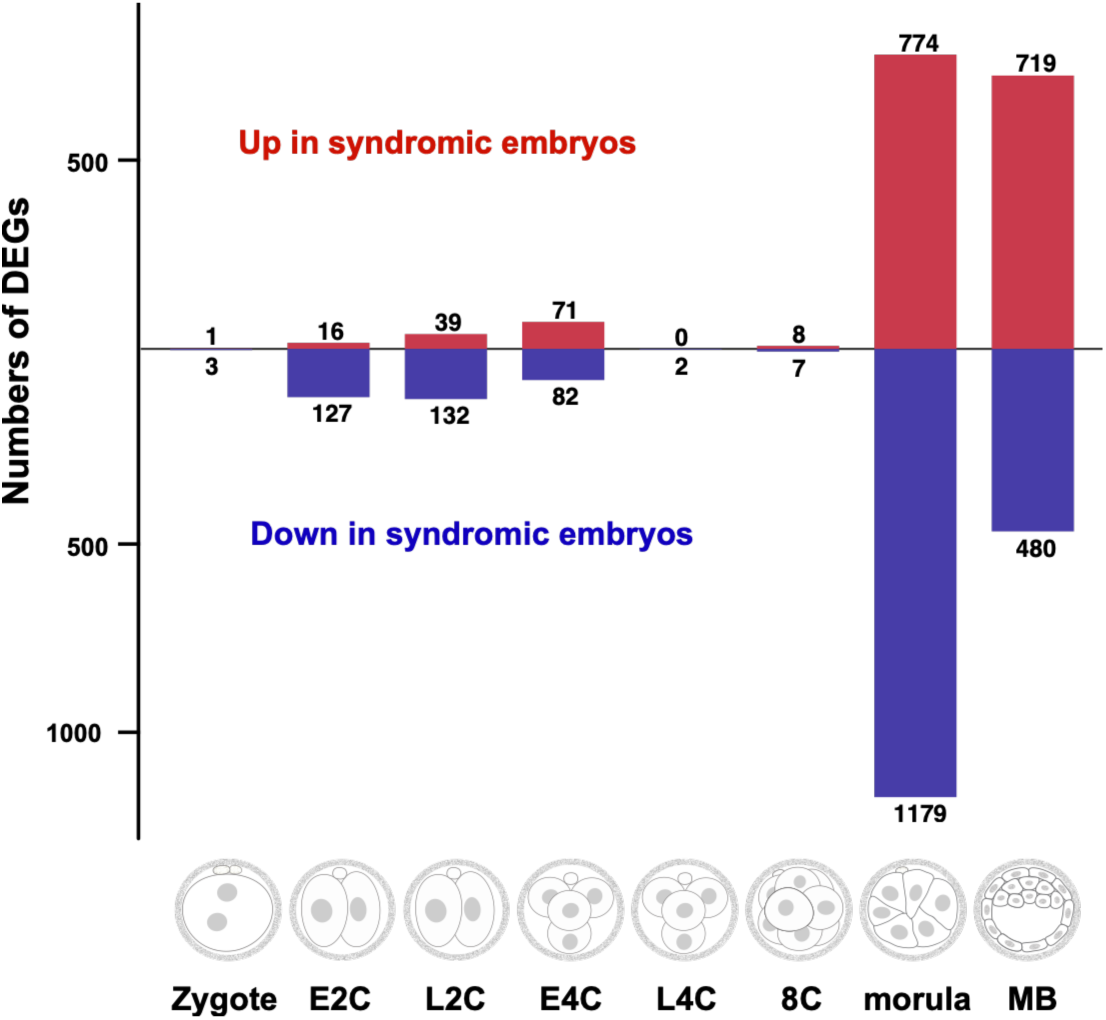
Numbers of DEGs between syndromic (Omd2 x BALB/c)F1 and control preimplantation embryos at each developmental stage. For each developmental stage, pairwise differential expression analyses were performed independently between syndromic (Omd2 x BALB/c)F1 embryos and each of the two control embryos independently: (i) non-syndromic (Omd2 x Omd2) embryos and (ii) non-syndromic (BALB/C x Omd2)F1 embryos. Genes were classified as DEGs only if they met the significance threshold (FDR < 0.1) in both pairwise comparisons, ensuring that the identified DEGs reflect the syndromic condition rather than strain-specific expression differences.

**Figure S9:**
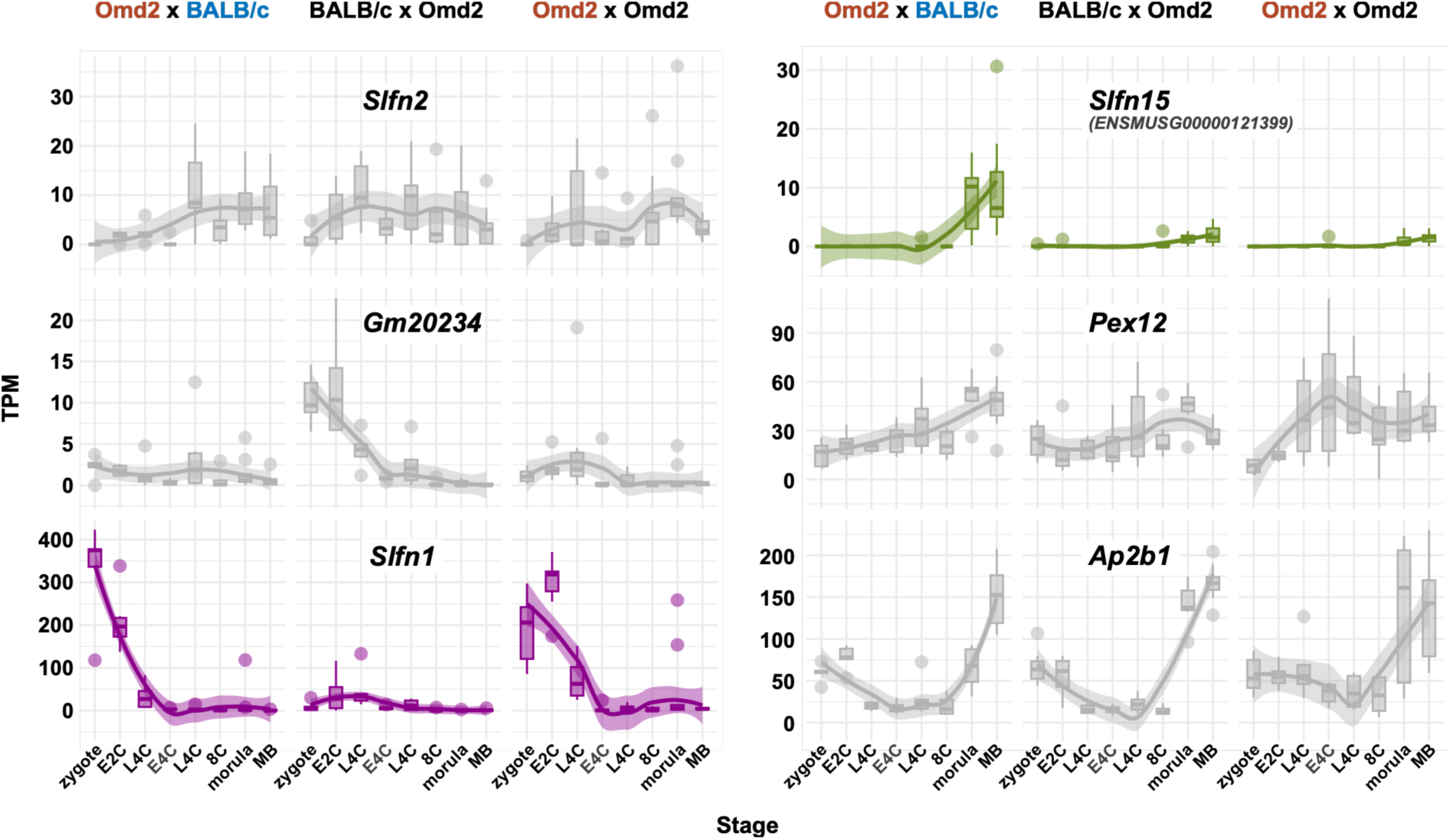
Expression profiles of *Om* region genes across preimplantation development. Transcripts per million (TPM) values at multiple developmental stages are shown for all *Om* region genes with a mean TPM≥10 in at least one condition. Data are presented for embryos from the three cross types: syndromic (Omd2 × BALB/c)F1, and non-syndromic control (Omd2 × Omd2) and (BALB/c × Omd2)F1 crosses.

**Figure S10:**
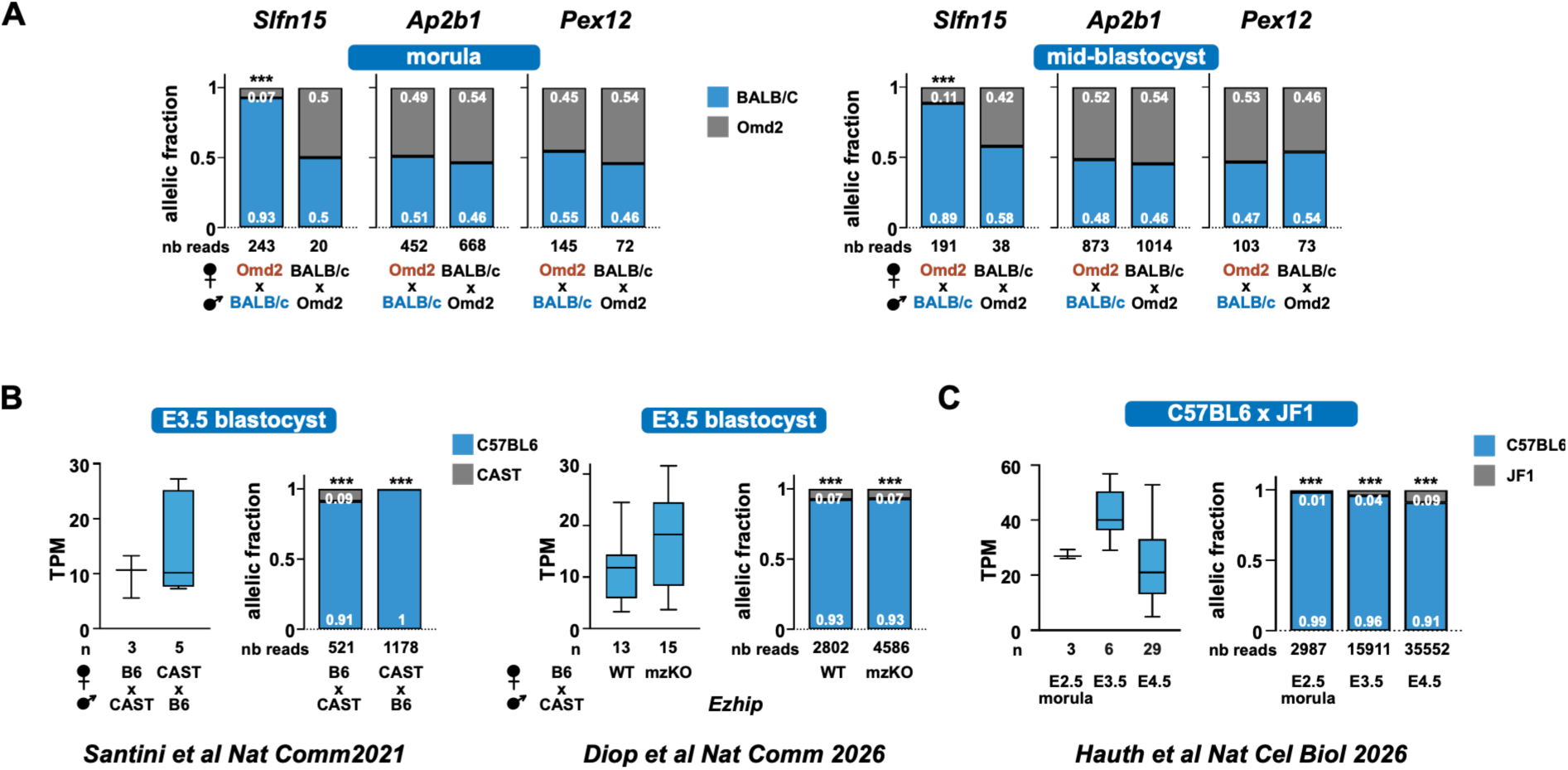
Allele-specific expression analysis of *Slfn15* in reciprocal hybrid embryos at preimplantation stages. **A.** Allele-specific *Slfn15* expression in (Omd2 × BALB/c)F1 and (BALB/c × Omd2)F1 embryos at the morula and mid-blastocyst stages. As controls, *Ap2b1* and *Pex12*, two genes located within the *Om* region and expressed at these stages, show no allelic bias in either cross confirming that allelic imbalance is specific to *Slfn15*. **B.** Allele-specific *Slfn15* expression in (C57BL/6 × CAST)F1 and (CAST × C57BL/6)F1 blastocysts at E3.5. **C.** Allele-specific *Slfn15* expression in (C57BL/6 × JF1)F1 embryos at morula (E2.5) and blastocyst (E3.5 and E4.5) stages. Note that both CAST (*Mus musculus castaneus*) and JF1 (*Mus musculus molossinus*) inbred strains lack the ETn/MusD retrotransposon insertion present in C57BL/6 and BALB/c incompatible strains. The number of informative reads from merged samples is indicated below each graph.

**Figure S11:**
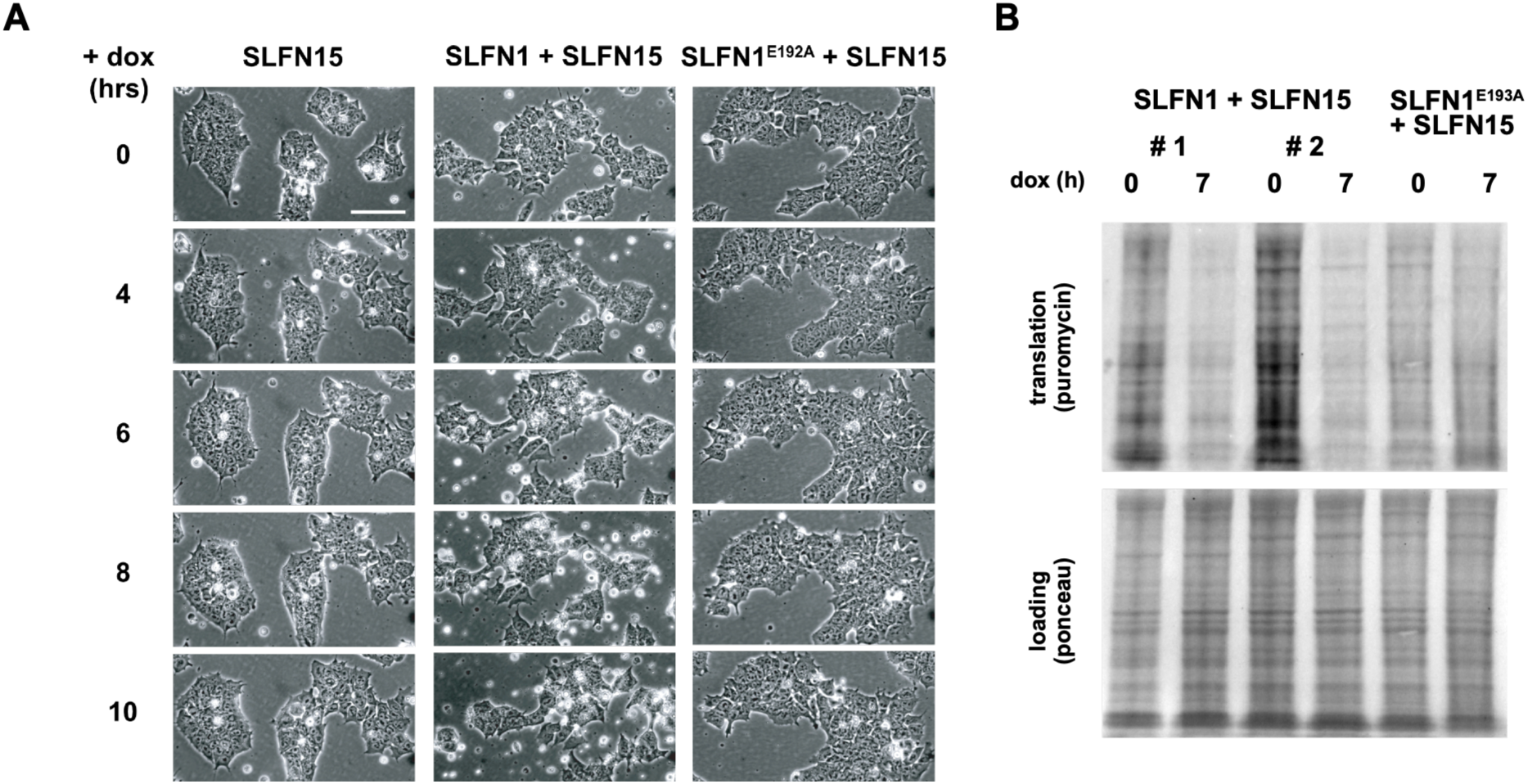
Morphological changes and inhibition of protein synthesis upon SLFN15 induction in ES cells expressing SLFN1 constitutively. **A.** Brigthfield and fluorescence images of KH2 ES cells expressing 3xFLAG-SLFN15 alone under dox-inducible control, or co-expressing either SLFN1-GFP or SLFN1^E193A^-GFP, taken at various timepoints following dox addition. After 6 hours of dox treatment, floating cells and debris begin to accumulate specifically in cells co-expressing 3xFLAG-SLFN15 and SLFN1-GFP, but not SLFN1^E193A^-GFP. Bar: 100µm. **B.** Assessment of global protein synthesis by ribopuromycylation assay in two independent SLFN1/SLFN15 clones and one SLFN1^E193A^/SLFN15 clone. Cells were treated or not with dox for 4 or 7 hours followed by puromycin incorporation for 10 min and analysed by immunoblotting with an anti-puromycin antibody.

**Figure S12:**
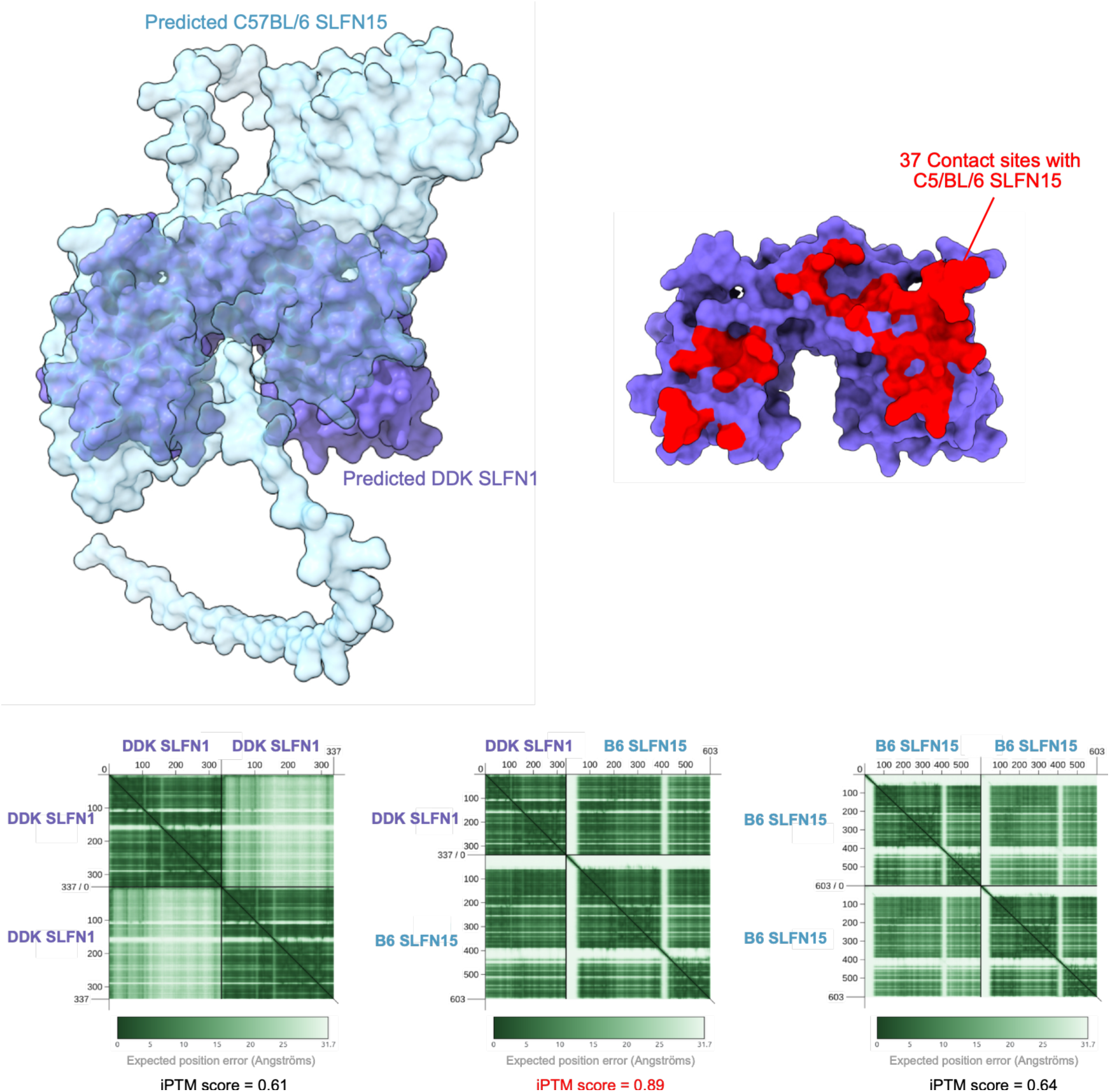
Alphafold3 prediction of SLFN1-SLF15 heterodimerisation. Alphafold3 was used to predict the structures of mouse SLFN1^DDK^ and SLFN15^B6^ monomers and to model their interaction. Residues on the SLFN1 protein surface predicted to engage in the dimerization interface with SLFN15 are highlighted in red. The interface predicted template modelling (ipTM) score for SLFN1-SLFN15 heterodimer exceeds 0.8, indicating a high-confidence prediction of protein-protein interaction The ipTM score predicted for SLFN1– SLFN15 heterodimerization is higher than those predicted for SLFN1–SLFN1 and SLFN15– SLFN15 homodimerization, suggesting preferential heterodimer formation.

**Supplementary Table S3:** *Slfn1* mRNA microinjection. The non-DDK strain used here is (C57Bl/6 x DBA2)F1.

| Microinjected RNA | Genotype of zygotes | Nb injected | Day 3 | Day 5 |  |  |
| --- | --- | --- | --- | --- | --- | --- |
|  |  |  | Viable (% injected) | Blastocyst (% viable) | Collapsed (% viable) | Degenerated (% viable) |
| *Slfn1<sup>DDK</sup>* | non-DDK x Omd2 | 113 | 81 (71.7) | 48 (59.2) | 16 (19.8) | 17 (21.0) |
|  | non-DDK x BALB/c | 115 | 78 (67.8) | 26 (33.3) | 7 (9.0) | 45 (57.7) |
| *Slfn1<sup>non-DDK</sup>* | non-DDK x Omd2 | 82 | 52 (63.4) | 39 (75.0) | 6 (11.5) | 7 (13.5) |
|  | non-DDK x BALB/c | 82 | 57 (69.5) | 18 (31.6) | 10 (17.5) | 29 (50.9) |

**Supplementary Table S4:**
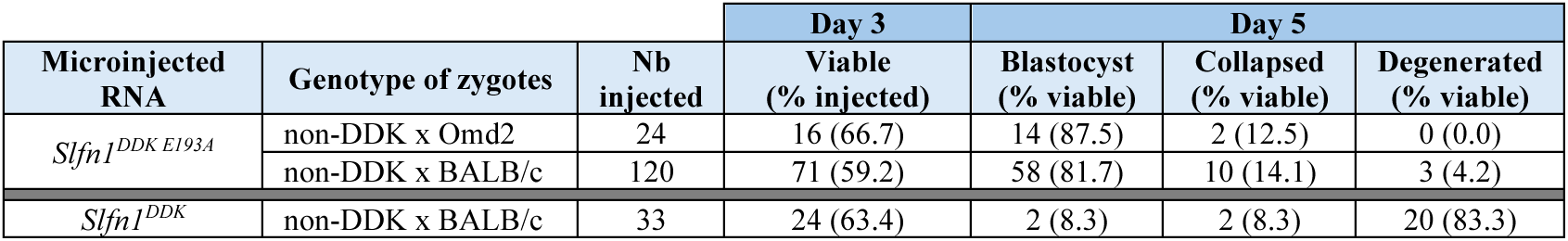
*Slfn1^E193A^* mRNA microinjection. The non-DDK strain used here is (C57Bl/6 x DBA2)F1.

**Supplementary Table S6:** *Slfn15* mRNA microinjection. The non-DDK strain used here is CD-1.

| Genotype of zygotes | Nb injected | Day 3 |  |  |  | Day 5 |  |  |
| --- | --- | --- | --- | --- | --- | --- | --- | --- |
|  |  | Lyzed (% injected) | 1C/2C (% injected) | 3C/4C (% injected) | Viable (% injected) | Blastocyst (% viable) | Collapsed (% viable) | Degenerated (% viable) |
| non-DDK x Omd2 | 37 | 5 (13.5) | 0 (0.0) | 3 (8.1) | 29 (78.4) | 23 (79.3) | 4 (13.8) | 2 (6.9) |
| Omd2 x Omd2 | 26 | 5 (19.2) | 19 (73.0) | 0 (0.0) | 2 (7.7) | 2 (100) | 0 (0.0) | 0 (0.0) |
| Omd2 x Omd2 | 55 | 2 (2.6) | 49 (89.0) | 1 (1.8) | 3 (5.5) | 0 (0.0) | 0 (0.0) | 3 (100) |
| ΔMERVL x Omd2 | 29 | 0 (0.0) | 6 (20.7) | 6 (20.7) | 17 (58.6) | 12 (70.6) | 3 (17.6) | 2 (11.8) |

**Supplementary Table S7:** *In vitro* development of syndromic embryos following CRISR/Cas9-mediated *Slfn15* deletion.

| Electroporation | Nb electroporated | Day 3 | Day 5 |  |  |
| --- | --- | --- | --- | --- | --- |
|  |  | Viable (% electroporated) | Blastocyst (% viable) | Collapsed (% viable) | Degenerated (% viable) |
| no | 38 | 35 (92.1) | 4 (11.4) | 0 (0.0) | 31 (88.6) |
| Without CRISPR/Cas components | 15 | 13 (86.7) | 1 (7.7) | 1 (7.7) | 11 (84.6) |
| Cas9 + *Slfn15* sgRNAs | 45 | 38 (84.4) | 29 (76.3) | 4 (10.5) | 5 (13.2) |

**Supplementary Table S8:**
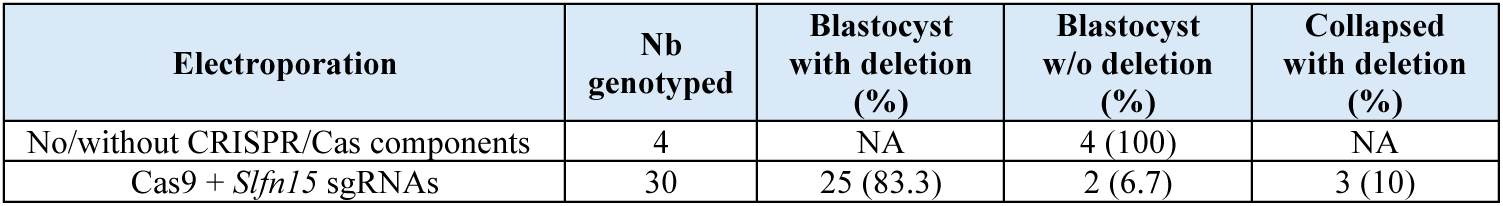
*Slfn15* allele modification frequency following CRISR/Cas9-mediated *Slfn15* deletion.

